# Bilayer thickness and transmembrane domain dynamics drive directional ER-Golgi traffic via Erv14/cornichon

**DOI:** 10.64898/2026.08.05.742935

**Authors:** Xiao-Han Li, Sangram Prusty, Nicolas Loyer, Maria Dolores Pejenaute Ochoa, Niamh Riley, Ellen Smith, Jens Januschke, Qiang Cui, Elizabeth A. Miller

## Abstract

Directionality of protein traffic during secretion is driven by sorting receptors that bind cargo in one compartment and release them in another. Erv14/cornichon family receptors mediate delivery from the endoplasmic reticulum (ER) to the Golgi. Erv14 traffics its client, Mid2, via a conserved interface formed by two transmembrane helices (TMHs). Molecular dynamics simulations of Erv14-Mid2 suggest that Erv14 TMHs undergo dynamic movement in response to both client binding and lipid bilayer thickness. Interaction between charged residues that flank the TMHs of Mid2 and Erv14 couple client binding to stabilization of the Erv14 export signal and thus engagement with the COPII coat. We term this mechanism Helix Engagement Induced Signal-mediated Traffic (HEIST) and suggest it as a general mechanism where changes in membrane biophysical properties drive directional transport.

## Introduction

One quarter of the eukaryote proteome comprises transmembrane proteins that perform diverse functions (*1*). Most of these proteins share common initial biogenesis steps: synthesis and folding in the endoplasmic reticulum (ER) and subsequent trafficking to the Golgi apparatus. ER-Golgi traffic largely uses COPII-coated transport intermediates (*2*). Cargo enrichment into COPII carriers occurs via cytoplasmic ER export signals that engage with Sec24, the cargo adaptor subunit of the COPII coat (*3, 4*). Cargo proteins can also use export receptors to connect indirectly to the coat (*5, 6*). Export receptors are transmembrane proteins that cycle between the ER and the Golgi, binding conditionally to their cargo clients in the ER and releasing them upon Golgi delivery thereby ensuring directional traffic (*5*).

The Erv14/cornichon family of ER export receptors are ubiquitous across eukaryotes (Fig. S1). The metazoan proteins are named after the *Drosophila melanogaster* family member, Cornichon (Cni), which plays a crucial role in TGFα signalling during development (*7, 8*). In humans, Cni family proteins comprise four paralogs, CNIH1-4, with CNIH1 and CNIH4 thought to retain function as cargo receptors (*9–11*), whereas CNIH2 and CNIH3 function as auxiliary subunits for AMPA receptors (*12–14*). Yeast Erv14 is the best characterized member of this family, responsible for export of diverse cargoes with the common property of long transmembrane helices (TMHs), a feature shared by many proteins that reside in the plasma membrane (*15–18*). Erv14 engages with Sec24 via an export signal in its cytoplasmic domain, which can augment independent export signals on cargoes (*15, 16, 18*).

Despite their conservation, it remains unclear how Erv14/Cni-family members conditionally recognize their client proteins to effect directional transport. The observation that Erv14 traffics cargo dependent on TMH length alone suggests that client recognition would require an unusual mechanism based on general hydrophobic interactions rather than a specific sequence motif (*17*). Such a binding mechanism could leverage biophysical effects of the lipid bilayer to drive reversible cargo engagement. To reveal the molecular basis of Erv14 function, we used an established client, Mid2, as a model to study how CNI-family proteins select and export their cargoes.

### Erv14 TMH1 and TMH4 drive cargo selection

We used AlphaFold2-multimer to predict structures of Erv14 in complex with the Mid2 TMH or a series of poly-leucine TMHs of increasing length (Fig. 1A). Reliable interactions were predicted for Erv14-Mid2_TMH_ and long-TMH cargoes (TMH >20aa) (Fig. S2A). Unbiased quantification of residue-specific contact probability suggests that the Mid2 TMH engages a membrane-embedded surface of Erv14 composed of TMH1 (to be referred to as H1) and TMH4 (H4). In contrast, cargo TMHs of 16- and 18-leucine repeats showed relatively low contact probability that was focused largely on H4 (Fig. S2B; 1A). Increasing TMH length increased contact probability with H1, closely resembling the Erv14-Mid2_TMH_ interaction (Fig. 1A). The extent of contact associated with increased TMH length recapitulated experiments measuring ER export of Mid2 chimeras, where Mid2-16L and Mid2-18L were ER-retained, and longer TMH-containing chimeras exported via Erv14 (*17*). Our modelling suggests Erv14 engages cargo via an interface comprising hydrophobic residues within H4 and a weakly hydrophobic/amphipathic surface in H1 (Fig. 1B,C). This putative interface is well-conserved, in contrast to higher variability of residues in TMH2 (H2) and TMH3 (H3) (Fig. 1D).

**Fig. 1.**
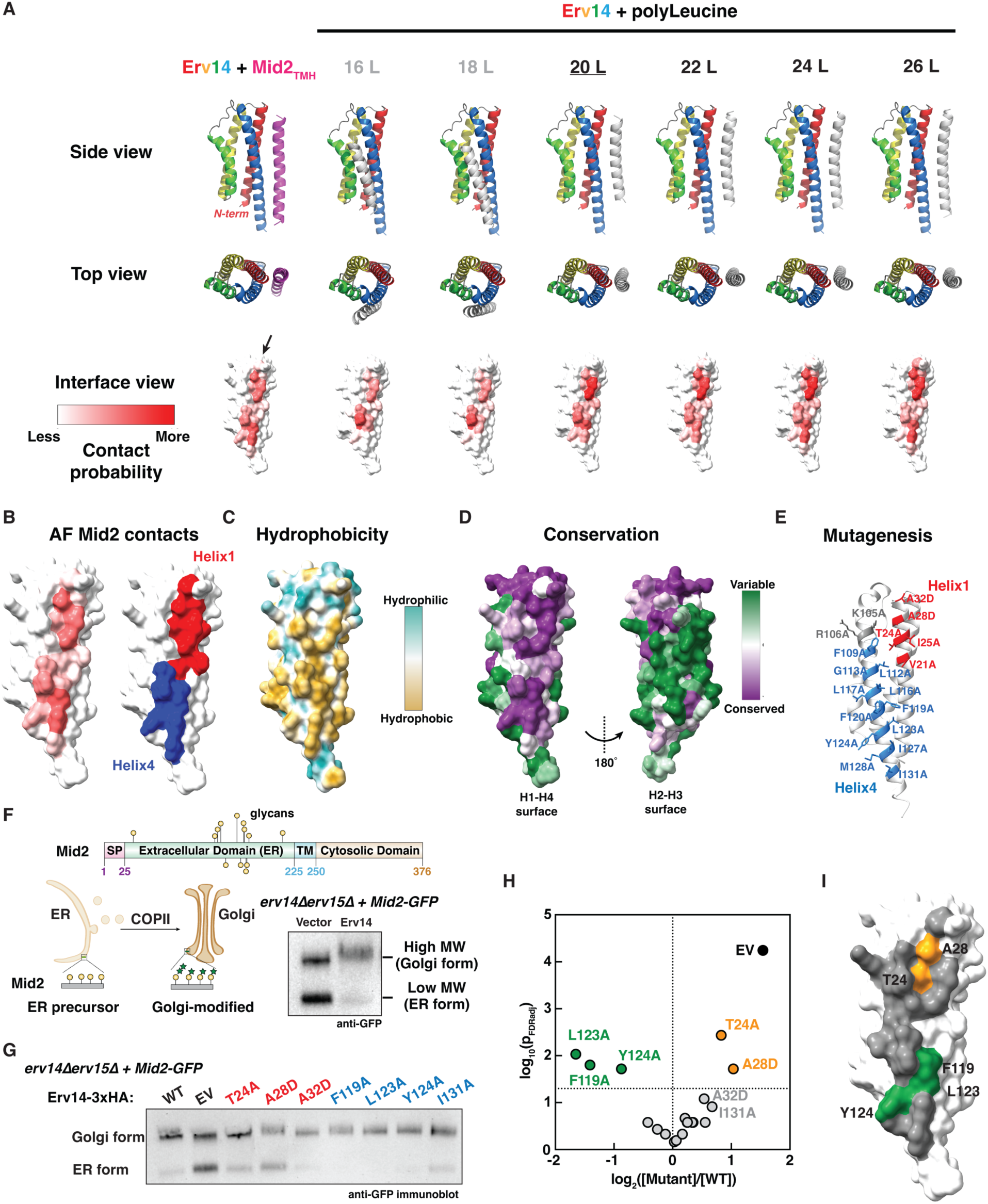
Erv14 Helix 1 and Helix 4 form a conserved interface for cargo recognition. (A) Top-rank predicted structures of yeast Erv14 (rainbow; H1-red; H2-yellow; H3-green; H4-blue) with Mid2_TMH_ (pink) and various lengths of polyleucine clients (grey), in top/side view cartoons and interface views rendered by the contact probability calculated from 30 predicted structures. (B) AlphaFold2 predicted contact interface (left) rendered by location of helix (right), with H1 in red and H4 in blue. (C) Erv14 predicted cargo-binding interface consists of a largely hydrophobic surface for TMH binding. (D) Conservation map of Erv14 (purple, well-conserved; green, poorly conserved) showing that residues on the H1-H4 interface are well-conserved in contrast to the H2-H3 surface. (E) Mutations on the H1-H4 interface of Erv14 chosen for mutagenesis. (F) Domain architecture of Mid2 (top) that shows predicted N-glycosylation (green) and O-glycosylation (yellow) sites. Scheme for immunoblot assay monitoring Mid2 glycosylation state as a proxy for ER/Golgi localization (bottom left), and example immunoblot that shows Golgi maturation of Mid2-GFP is enhanced by Erv14 (bottom right). (G) Immunoblot of Mid2 maturation expressed in the context of distinct Erv14 mutants that showed various LoF, GoF, or neutral phenotypes. (H) Volcano plot showing quantification of Mid2 trafficking in the presence of Erv14 mutants. LoF and GoF mutants are colored in orange or green, respectively, neutral mutants are colored in grey, and empty vector control (EV) in black. Each point represents one mutant, measured across multiple repeats (n=4-5) with adjusted p value determined by one-sample t-test against null hypothetical mean 0, followed by Benjamini–Hochberg correction with false discovery rate (FDR) at 0.05. The significance cutoff at p_FDRadj_ = 0.05 is shown as a horizontal line at log₁₀(p_FDRadj_) = 1.3. (I) Location of LoF and GoF mutants on the H1-H4 interface.

We tested the functionality of the Erv14 H1/H4 interface using site-directed mutagenesis (Fig. 1E). Plasmids bearing mutations in individual amino acids were introduced into a strain expressing Mid2-GFP and deleted for both *ERV14* and its partially redundant paralog, *ERV15*. Stable expression of each mutant was confirmed by immunoblot (Fig. S3A). We measured Mid2 traffic by quantifying the proportion of low molecular weight ER-localized Mid2 (Fig. 1F upper panel) (*17, 18*). In *erv14Δerv15Δ* cells, Mid2 leaves the ER slowly, resulting in a significant population of ER-retained protein. Introduction of wild type Erv14 significantly decreased this ER-retained pool (Fig. 1F). Quantification of the proportion of ER-retained Mid2-GFP across multiple independent experiments revealed five point mutations that resulted in altered Mid2 traffic (Fig. 1G-H; Fig. S3B). Two mutations in H1 (T24A, A28D) significantly impaired ER export (loss-of-function, LoF) of Mid2 (Fig. 1G-I), suggesting an active role in cargo-binding.

Surprisingly, three substitutions within H4 (F119A, L123A, Y124A) appeared to accelerate Mid2 export (gain-of-function, GoF; Fig. 1G-I), perhaps by contributing to helix dynamics that would favour either Mid2 binding or ER export.

### Erv14 TMH dynamics couple bilayer thickness and cargo binding

Our model of directional traffic driven by biophysical effects of the lipid bilayer derives in part from established differences in bilayer thickness between the ER and downstream organelles (*19–21*). Within the ER, long TMHs are subject to hydrophobic mismatch, which can influence membrane protein behaviour and interactions (*22–24*). We thus sought to test how the local lipid environment influences Erv14 and Mid2 behaviour. Using the best-ranked structural predictions of Erv14, Mid2_TMH_, and the Erv14-Mid2_TMH_ complex as starting models, we performed all-atom MD simulations in two different single-component lipid bilayers: DRPC (thin, mimicking ER) and DSPC (thick, mimicking Golgi)(*25*). In thick DSPC membranes, Erv14 induced local thinning of the bilayer, whereas in thin DRPC bilayers, Erv14 increased local bilayer thickness (Fig. 2A, Fig. S4A). In the context of Erv14-Mid2_TMH_, these effects on bilayer thickness were amplified compared to Erv14 alone (Fig. S4B). The data suggest that Erv14 modulates local bilayer thickness, and can buffer bilayer thickness to a specific range, possibly facilitating cargo binding.

**Fig. 2.**
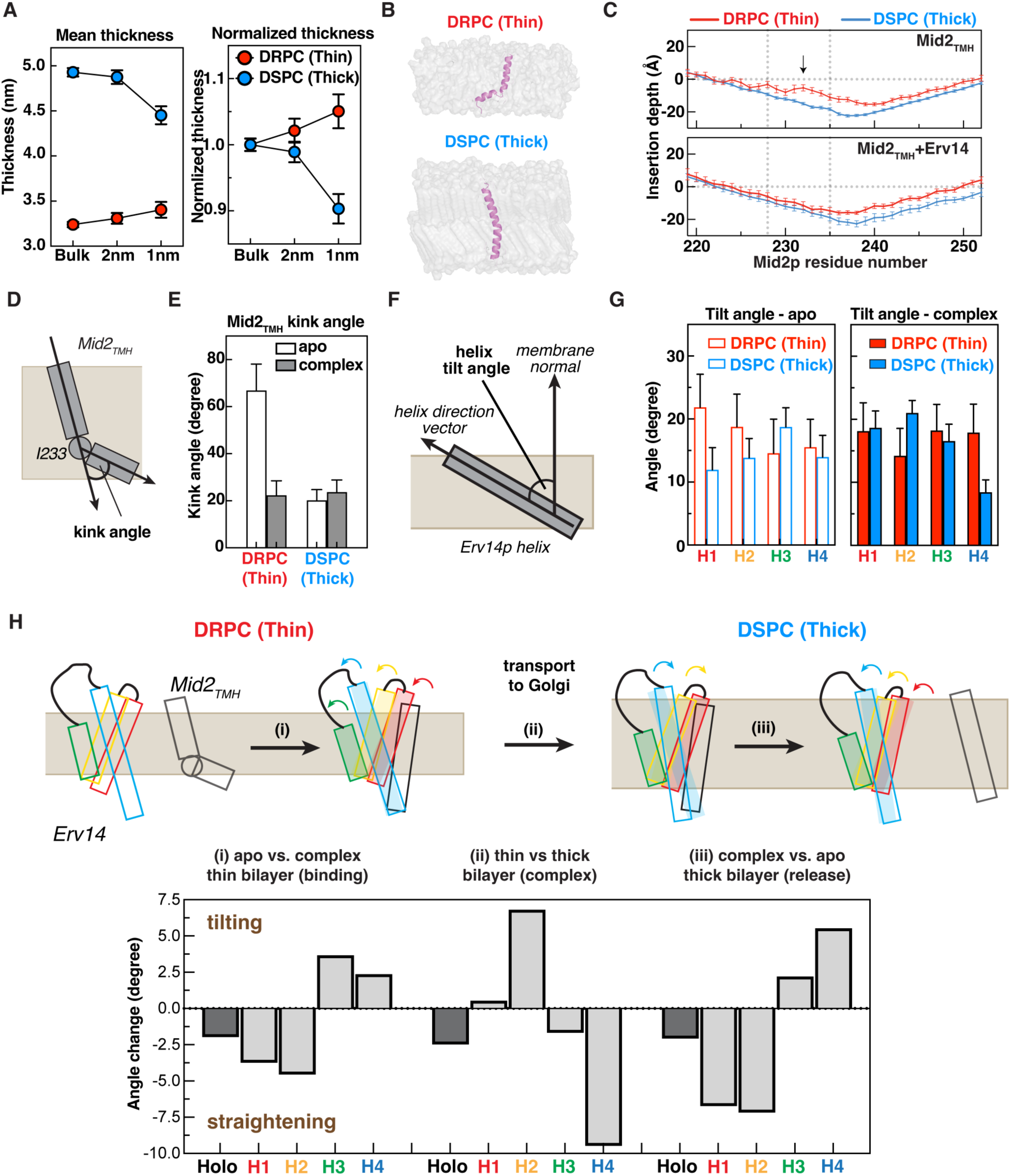
Erv14 modulates bilayer thickness and dynamically engages cargo via helix tilting. (A) Bilayer thickness (left) and normalized change in thickness (right) of different bilayers as a function of distance from the surface of Erv14 as measured by MD simulation. (B) Snapshots of representative conformations of Mid2_TMH_ (magenta) in DRPC/DSPC bilayer, showing the helix kink. (C) Residue-specific bilayer insertion depth of Mid2_TMH_ without (top) or with (bottom) Erv14 engagement in thin (red) or thick (blue) lipid bilayers; the kinked helical region 228-235 is marked by dashed lines and arrow. (D) Scheme depicting definition of helix kink angle in Mid2_TMH_. (E) Mid2_TMH_ helix kink angle in the presence (solid) or absence (clear) of Erv14 in different bilayer thickness. (F) Scheme depicting the definition of helix tilt angle. (G) Individual helix tilt angles for Erv14 TMHs in different bilayers and without (left) or with (right) Mid2_TMH_ engagement. (H) Scheme showing comparisons that model distinct steps of Mid2 traffic (upper panel). Helix tilt angle changes for Erv14 as a whole (lower panel; dark grey), or individual TMHs (lower panel; light grey) reveal how Erv14 changes under different conditions. Movement of each TMH in each step is indicated in the cartoon (upper panel) by outlined rectangles (current state), shaded rectangles (previous state), and arrows (direction of movement).

When modelling the behaviour of Mid2_TMH_ alone, we observed distinct conformations in thin versus thick bilayers, likely reflecting the need to accommodate hydrophobic mismatch in the thin DRPC bilayer (Fig. 2B). In DRPC, Mid2_TMH_ adopted a kinked structure with the TMH broken around residue I233, reflected by shallow insertion depth across residues 228-235 (Fig. 2C). In the Erv14-Mid2_TMH_ complex the kink was rectified, with deeper insertion for residues 228-235 and a continuous increase in insertion depth across the TMH (Fig. 2C; Fig. S5A). In contrast, in thick DSPC bilayers, Mid2_TMH_ showed uniform increased insertion depth across the TMH and was not altered by Erv14 (Fig. 2B-C; Fig. S5A). We further quantified TMH behaviour by measuring the kink angle centred at residue I233 (Fig. 2D-E), revealing resolution of the large kink angle by Erv14 binding in DRPC, or by insertion in DSPC (Fig. 2E). This behaviour of the Mid2 TMH led us to reconsider the GoF mutations in Erv14-H4 (Fig. 1F-H). We leveraged our MD simulations to quantify residue contacts between Erv14-H4 and Mid2_TMH_, revealing four Erv14 residues (F119, F120, L123, Y124) that bind to the kink region of Mid2_TMH_ in both DRPC and DSPC (Fig. S5B). This H4 tetrad may function to “drag” the Mid2 helix into the bilayer, contributing to kink rectification and buffering against hydrophobic mismatch. The three GoF mutations, F119A, L123A, Y124A, that sit in close proximity to the bulky hydrophobic side chains near the Mid2 kink (Fig. S5C), are well placed to reduce the energy costs of Mid2 engagement with Erv14 and therefore accelerate binding and export of Mid2.

We reasoned that Erv14 might also undergo dynamic TMH tilting in different bilayers. Indeed, when modelled in the apo state, H1 and H2 showed larger tilt angles in thin bilayers versus thick (Fig. 2F-G). In contrast, H3 and H4 tilt angles were relatively unchanged across bilayer conditions, with only H3 showing a slightly larger tilt angle in DSPC versus DRPC (Fig. 2G, left panel). In the Erv14-Mid2_TMH_ complex, H1 and H3 remained unchanged across bilayer conditions whereas H2 and H4 changed orientation (Fig. 2G, right panel). The distinct behaviour of individual helices suggests a concerted conformational change and hence a mechanism for conditional client engagement. We therefore measured changes in TMH behaviour during progression from (i) Erv14 monomer to Erv14-Mid2_TMH_ complex in DRPC (modelling client binding in ER), (ii) Erv14-Mid2_TMH_ complex in DRPC versus DSPC (modelling delivery to Golgi), and (iii) Erv14 complex versus monomer in DSPC (modelling client release). Quantifying the change in tilt angles of holo-Erv14 revealed straightening at each step (Fig. 2H). First, in the thin bilayer, Mid2_TMH_ engagement and kink rectification straightens Erv14 to accommodate the long Mid2-TMH and locally thicken the bilayer. Delivery to the thick bilayer of the Golgi requires further straightening to prevent hydrophobic mismatch, but the degree of mismatch is buffered by the long Mid2-TMH. Finally, upon Mid2 release, straightening even further maximizes the hydrophobic nature of the Erv14-TMHs alone. Interestingly, individual Erv14-TMHs seemed to undergo distinct bilayer-dependent tilting behaviour during this progression (Fig. 2H). Specifically, in the thin-to-thick transition H2 and H4 undergo significant changes in tilt angle whereas H1 and H3 remain static (Fig. 2H). Our interpretation of this behaviour is that H2/H4 movement upon delivery to the Golgi could alter the H1/H4 interface, favouring eviction of the cargo. We thus propose that bilayer-sensitive dynamics of specific Erv14 helices couple membrane thickness to conditional cargo binding.

### Coupling of helix dynamics and ER-export is conserved in cornichon

Conservation of the H1/H4 interface led us to test if other cornichon family proteins could use the same mechanism for ER export. We turned to the fly model, where the *cornichon* (*cni*) mutant causes oogenesis defects resulting from defective GURKEN (Grk, fly TGFa) signaling (*7, 26*). We modelled the structure of Cni in complex with Grk_TMH_, comparing the best-ranked model to the Erv14-Mid2_TMH_ model, and chose three sets of mutations that likely perturb Cni in a similar manner as Erv14 (Fig. 3A). LoF mutations to the H1 interface should impair cargo selection (*cni^I25A,I29A^*); mutations to the H4 GoF tetrad should accelerate export (*cni^I134A,Y135A^*); mutations to H3-H4 cytoplasmic COPII binding loop should block export by impairing Sec24 engagement (*cni^104-108A^*). We first examined recruitment of Cni at ER exit sites (ERES) marked by Sec16::GFP in egg chambers (Fig. 3B), observing that both LoF and COPII binding mutants lost ERES localization in nurse cells, whereas the GoF mutant increased its abundance at ERES (Fig. 3B-C). Together the LoF and GoF localization phenotypes are consistent with a model of cargo engagement directly influencing recruitment to COPII assembly sites.

**Fig. 3.**
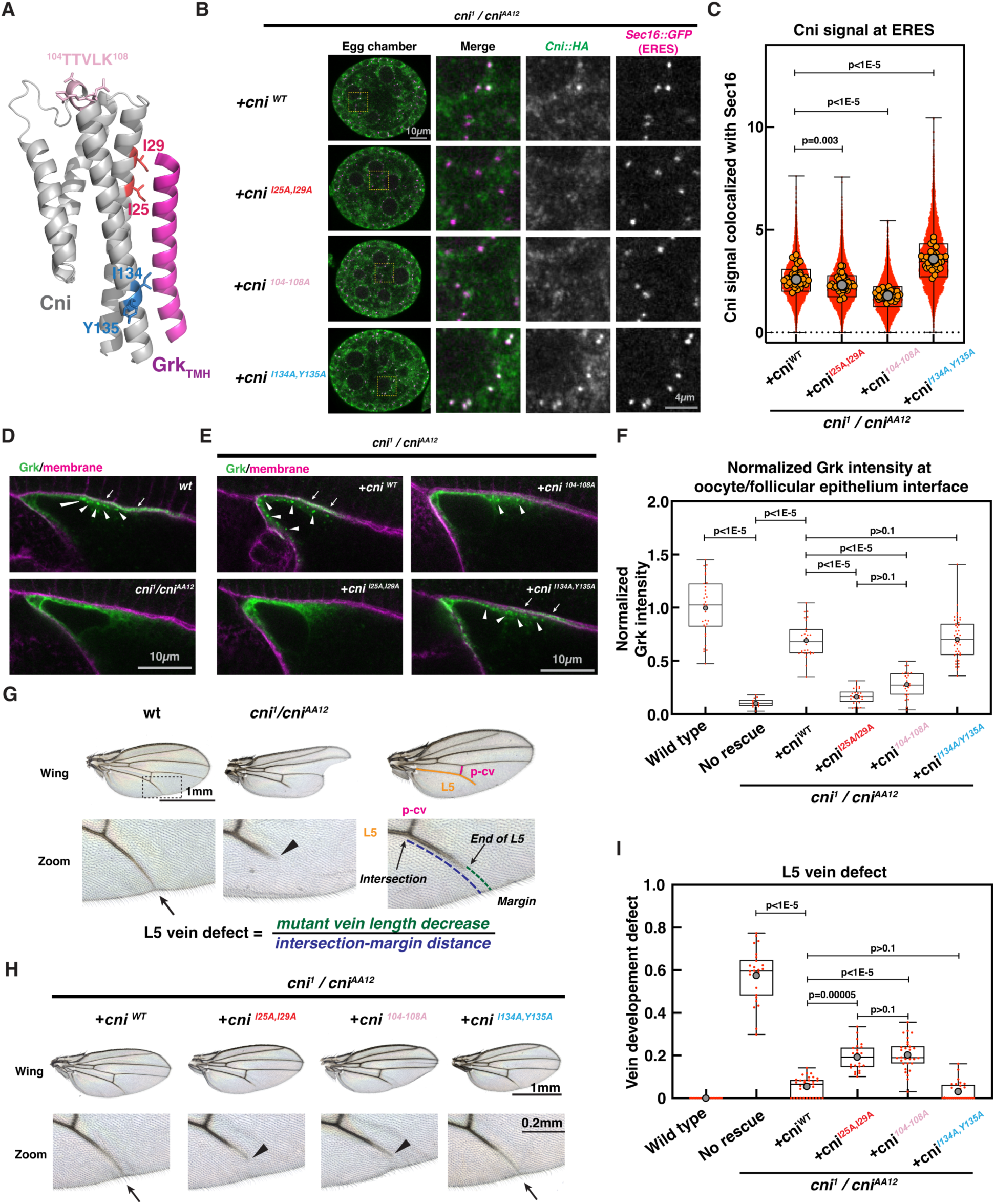
Drosophila cornichon (Cni) engages clients by a conserved mechanism. (A) Predicted structure of Drosophila Cni (grey)-Grk_TMH_ (magenta) complex, highlighting mutants corresponding to H1 LoF (red), H4 GoF (blue), and COPII binding (pink). (B) Immunofluorescence imaging of Drosophila egg chambers showing colocalization of HA-tagged Cni variants (green) and ERES marked by Sec16::GFP (magenta) in nurse cells. Regions in yellow squares are enlarged to visualize signals from different channels. (C) Quantification of Cni signal at ERES marked by Sec16::GFP. Red dots represent individual Sec16-positive puncta, orange circles represent mean of each individual egg chamber, grey dot represents mean of entire dataset. (n=5-6, statistical significance determined by ordinary one-way ANOVA followed by post-hoc Dunnett’s test against Cni^WT^). (D) Immunofluorescence imaging of Drosophila oocyte showing Grk (green) localizes to punctate structures (arrow heads) and plasma membrane (magenta; arrows) in *wild type* flies but is largely intracellular in a *cni^1^/cni^AA12^* line. (E) Immunofluorescence imaging of Drosophila oocytes expressing the indicated Cni variants in a *cni^1^/cni^AA12^* background, highlighting intracellular puncta (arrowheads) and membrane co-localization (arrows). (F) Quantification of normalized Grk intensity at the membrane interface is shown in the bottom panel (wild type n=31; no rescue n=14; cni^WT^ n=28; *cni^I25A,I29A^*n=26; *cni^I134A,Y135A^* n=43; cni^104-108A^ n= 25; statistical significance determined by ordinary one-way ANOVA followed by post-hoc Tukey’s test). (G) Adult wings from wild type or *cni^1^/cni^AA12^* lines (top) and higher magnification view of L5 vein area (bottom). Arrows and arrowheads indicate wildtype and impaired vein development, respectively. Wing on right shows measurement scheme for L5 from the intersection of posterior cross-vein (p-cv) to the wing margin. The wing development defect is quantified as the ratio between the decrease in L5 vein length in mutants over projected L5/p-cv intersection to wing margin distance. (H) L5 wing phenotypes of different mutants expressed in *cni^1^/cni^AA12^*line. (I) Quantification of L5 vein defects in wild type, *cni^1^/cni^AA12^*, and the indicated mutants in *cni^1^/cni^AA12^* line (wild type n=20; no rescue n=26; cni^WT^ n=33; *cni^I25A,I29A^*n=25; *cni^I134A,Y135A^* n=29; cni^104-108A^ n=30; statistical significance determined by Kruskal-Wallis test followed by post-hoc Dunn’s test).

We then tested functionality of *cni* mutants in Cni-defective *cni^1^/cni^12AA^* transheterozygote females (*7*), looking first at Grk localization in oocytes. In WT oocytes, Grk showed significant cortical localization (marked with phalloidin) at the interface with neighbouring follicular cells, indicative of delivery to the cell surface. In contrast, in the *cni^1^/cni^12AA^* line, Grk was largely diffuse and failed to co-localize with the cortical marker (Fig. 3D). Introduction of either WT Cni or the GoF H4 mutant rescued this phenotype, whereas the COPII-binding and H1 interface mutants failed to rescue (Fig. 3E-F), consistent with conserved mechanisms mediating function across family members.

In addition to defective Grk localization in oocytes, *cni^1^/cni^12AA^*adults showed altered wing shape and disrupted L5 vein (Fig. 3G) (*8*). Introduction of WT Cni into *cni^1^/cni^12AA^* mutants largely restored the wing phenotype and L5 vein growth; LoF mutations in the H1 interface and COPII-binding loop failed to rescue L5 vein defects; the GoF H4 mutant showed a similar rescue to *cni*^WT^ (Fig. 3H-I), mirroring our observation in rescue of Grk localization at the membrane (Fig. 3D-F). Our results suggest a evolutionarily conserved mechanism for cornichon proteins in driving ER-export.

### Erv14 cytoplasmic loops couple cargo binding to ER export signal engagement

The Cni ERES localization phenotypes and Erv14 dynamics in different bilayers led us to consider how TMH movement might contribute to directional traffic by coupling recruitment of Sec24 to the formation of the Erv14-Mid2 complex. Our MD simulations revealed significant differences between insertion depth of residues contained within two Erv14 cytoplasmic loops, suggesting functional importance corresponding to bilayer thickness (Fig. 4A). The H3-H4 loop contains a known export signal that is thought to engage the Sec24 “D-site” (*15, 16, 18*). The H1-H2 loop contains highly conserved negatively charged residues previously shown to be important for plant cornichon function (*27*) (Fig. 4A). Mutagenesis across the H1-H2 loop confirmed its contribution to Mid2 traffic, with the strongest effect observed for residues closest to H2; mutating the most conserved residues in the H3-H4 loop TEIFR shows similar LoF as the well-characterized IFRTL signal, highlighting the importance of the ^97^IFR^99^ motif in COPII binding (Fig. 4B; Fig. S3C). Stability of each mutant was confirmed by immunoblot (Fig. S3B). AlphaFold2 predicted a high-confidence structure of the Erv14-Mid2-Sec24 complex, revealing a possible multipartite interface (Fig. 4C). The Sec24 D-site pocket was in close proximity to both the ^97^IFR^99^ motif and the H1-H2 loop (Fig. 4C inset). Moreover, negative residues in the H1-H2 loop of Erv14 were positioned to engage positive residues in the Mid2 cytoplasmic domain, which in turn flank a series of acidic residues close to the Sec24 B-site. Together, the structural model suggested a mechanism of cooperative engagement of the Erv14-Mid2 complex with Sec24: Erv14 binds to Sec24 via the H3-H4 loop (IFR motif); the Erv14 H1-H2 loop engages both the Sec24 D-site and positively charged residues in Mid2; this interaction positions the Mid2 acidic residues in an orientation that permits binding by the Sec24 B-site, thereby creating a robust combinatorial interaction. We tested the importance of the Mid2 charged residues, finding that only the combination of alanine substitution at both positively and negatively charged sites significantly impaired export (Fig. 4D).

**Fig. 4.**
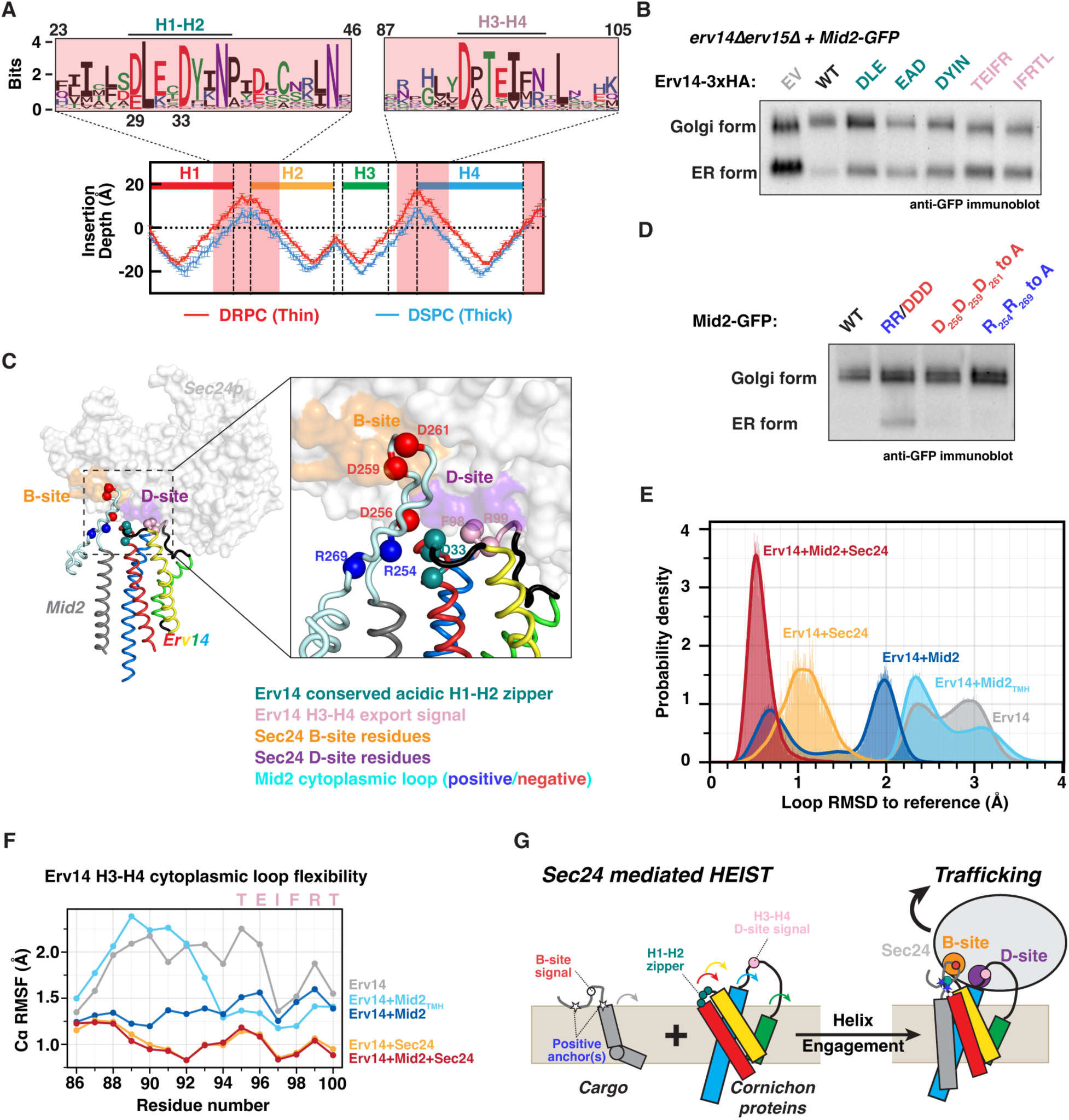
Erv14 leverages helix engagement to expose multivalent Sec24 binding signals. (A) Residue-specific insertion depth of Erv14 in different lipid bilayers as calculated for the Erv14-Mid2_TMH_ complex. Dashed lines and coloured bars on top mark the boundaries of each helix. Shaded area corresponds to robust exposed residues in DRPC. Residue conservation across the exposed regions are shown as sequence logos (top), with conserved H1-H2 (teal) and H3-H4 (pink) regions highlighted. (B) Immunoblot analysis of Mid2-GFP trafficking for H1-H2 and H3-H4 mutants. (C) Predicted structure of Erv14-Mid2-Sec24, highlighting the key residues as spheres: Erv14 H1-H2 zipper (teal), Erv14 H3-H4 export signal (pink), Mid2 arginine pair (blue), Mid2 triple negative signal (red). Sec24 B- and D-sites are shaded orange and purple, respectively. (D) Immunoblot analysis of Mid2-GFP trafficking for indicated Mid2 cytoplasmic mutants. (E) MD-simulated RMSD distribution of the Erv14 H3-H4 loop in complexes of Erv14 (grey), Erv14-Mid2_TMH_ (cyan), Erv14-Mid2 (blue), Erv14-Sec24 (orange) and Erv14-Mid2-Sec24 (red), with Erv14-Mid2-Sec24 as reference structure, showing stepwise convergence of Erv14 export signal loop towards a three-member complex. (F) MD-simulated residue-specific RMSF for the Erv14 H3-H4 loop in the indicated complexes, showing the stepwise decrease in flexibility as Mid2 and Sec24 engage. (G) Proposed HEIST mechanism for Sec24-mediated cargo trafficking.

To understand how these combinatorial interactions couple Sec24 engagement to cargo selection, we used MD simulations to examine the dynamics of the Erv14 H3-H4 loop, which contains the IFR export signal (Fig. 4E). We first analysed root mean square displacements (RMSDs) for simulations of various subcomplexes in the thin bilayer mimicking the ER environment. RMSD reports on structural deviations of each subcomplex compared to the full tripartite complex as a reference. For Erv14 alone, the H3-H4 loop showed two RMSD populations centred at 2.3 Å and 3.0 Å (Fig. 4E). Binding of Erv14 to Mid2_TMH_ increased the relative population of lower RMSD state at 2.3 Å; adding the Mid2 cytoplasmic domain (Erv14-Mid2) further reduced the RMSD, with two populations centred at 0.7 Å (close to the reference) and 2.0 Å (Fig. 4E). Interestingly, RMSD of the Erv14-Sec24 complex was intermediate between the two populations of the Erv14-Mid2 complex, suggesting that the presence of Mid2 creates a population of Erv14 that is more stable or capable of binding Sec24. To gain further clarity on contributions of individual residues to loop dynamics, we calculated residue-specific root mean square fluctuation (RMSF) values for the two cytoplasmic Erv14 loops, reporting residue-level flexibility. The H3-H4 loop was most flexible in Erv14 alone, with stepwise changes as elements of the complex were added (Fig. 4F). The Erv14-Mid2_TMH_ complex stabilized residues surrounding the IFR signal; in the Erv14-Mid2 complex that included the Mid2 cytoplasmic domain, further stability across the entire loop was apparent. Addition of Sec24 further reduced residue-level fluctuations, consistent with this state being the most stable structure (Fig. 4F). In contrast, both the RMSD and flexibility of the H1-H2 Erv14 loop changed only minimally across the different complexes (Fig. S6), suggesting this loop functions more as a structural bridge that links the various structural components.

### HEIST: a conserved mechanism for ER-Golgi traffic of membrane proteins

Together, our integrated approach using structural modelling, MD simulation and functional analysis in yeast and flies suggest a conserved mechanism for Erv14-mediated ER export (Fig. 4G). Capture of a long TMH at the Erv14 H1/H4 interface partially rectifies client TMH hydrophobic mismatch and locally thickens the lipid bilayer. Engagement of the Erv14 acidic H1-H2 loop with positively charged residues in the client’s cytoplasmic domain stabilizes the complex and induces cooperative binding to Sec24. Upon delivery to the thick bilayer of the Golgi, conformational changes in Erv14 TMHs prise apart the H1/H4 interface, releasing the client TMH and retracting the Sec24-engaged residues towards the lipid surface. We term this mechanism HEIST, for Helix Engagement Induced Signal-mediated Traffic, and propose that similar principles may operate at other trafficking events where membrane changes might impact TMH behaviour to drive directional transport.

Three key features in Mid2 contribute to its Erv14-mediated HEIST: (1) a long TMH that causes hydrophobic mismatching in the ER; (2) positively charged cytoplasmic residues flanking the TMH; (3) negative charges in the cytoplasmic domain as a candidate Sec24 B-site signal. These general features are conserved across potential clients. A long TMH is established as a feature driving Erv14 engagement (*17*). Enrichment of positively charged residues on the cytoplasmic face of transmembrane proteins is a general feature known as the “positive-inside” rule (*28, 29*). We propose that the conserved H1-H2 loop residues in cornichon proteins act as an electrostatic “zipper” to stabilize Erv14-client interaction and position putative ER export signals (Fig. 4).

Predictions of Erv14-client-Sec24 complexes for other cornichon cargoes (yeast Axl2p; fly Grk and Notch) suggest conserved TMH engagement via H1-H4 and positive charge-zipper interactions, whereas client cytoplasmic domains that might contribute to Sec24 engagement were more variable (Fig. S7). All clients had a combination of negatively charged residues and bulky hydrophobic residues that could feasibly drive B-site interactions, albeit in distinct arrangements. Such variability suggests plasticity in how the Sec24 B-site engages with different clients, thereby increasing diversity of cargos that can be exported via HEIST.

Polytopic transmembrane proteins are the most abundant class of Erv14 clients in yeast. Structural analysis of an Erv14-Qdr2 complex independently identified the H1/H4 surface as the cargo-facing interface and revealed extensive lipid-mediated remodeling around this long-TM polytopic cargo [Tunyi *et al.,* 2026]. The observation that one or more Erv14 molecules likely buffer hydrophobic mismatch and position cargo export motifs for Sec24 engagement is consistent with the central HEIST principle, but suggests that receptor stoichiometry and fine architecture at the interface may vary with cargo topology. To further understand the HEIST model for a polytopic membrane protein with a large cytoplasmic domain and known export signal, we performed structural predictions of Yor1-Erv14-Sec24, where Yor1 is an ABC transporter that traffics via Erv14 and the Sec24 B-site (*3, 18, 30, 31*). The model predicts that the Erv14 H1-H4 interface engages Yor1 via a short region of TMH10, centered around the GoF triad. This relatively small interface is supplemented by a Yor1p cytoplasmic loop that contains bulky hydrophobic residues flanked by positive charge clusters that lies close to the Erv14 H1-H2 zipper (Fig. S8). This model would be consistent with further stabilization of Erv14-Yor1 interface via lipids, as seen in the Erv14-Qdr2 structure [Tunyi *et al.,* 2026]. Importantly, the tripartite model predicts that the known ER-export signal for Yor1 (^71^DIE^73^) is positioned adjacent to the B-site, consistent with HEIST-mediated export of Yor1. Taken together, we propose that HEIST serves as a robust mechanism to integrate weak sorting signals with biophysical features of transmembrane proteins and thus achieve robustness and efficiency in protein trafficking.

## Acknowledgments

The authors used ChatGPT 5.4 (OpenAI) to assist with drafting and debugging Python scripts used for bioinformatics and visualizations. The tool was not used to generate or interpret scientific conclusions. All code was inspected, validated against expected outputs, and edited by the authors, who take full responsibility for the final analyses and results. Computational resources provided by the Boston University Shared Computing Cluster, which is administered by Boston University Research Computing Services (www.bu.edu/tech/support/research/), are greatly appreciated. Part of the computation also used the ACES GPU cluster at Texas A & M University through allocation BIO260037 from the <u>Advanced Cyberinfrastructure Coordination Ecosystem: Services & Support</u> (ACCESS) program, which is supported by U.S. National Science Foundation grants #2138259, #2138286, #2138307, #2137603, and #2138296. The authors thank the Data Analysis Group and DTS Research Computing HPC service at the University of Dundee for providing computational resource for structural predictions. The authors thank R. Hegde, S. Munro, M. Lee, H. Wu for carefully reading the manuscript and helpful suggestions.

## Author contributions

Conceptualization: XHL, QC, EAM; Methodology: XHL, SP, NL, MDPO, JJ, QC, EAM; Investigation: XHL, SP, NL, MDPO, ES, NR; Visualization: XHL, SP, NL, MDPO, JJ, QC, EAM; Funding acquisition: QC, EAM; Project administration: QC, EAM; Supervision: JJ, QC, EAM; Writing – original draft: XHL, EAM; Writing – review & editing: XHL, SP, NL, MDPO, JJ, QC, EAM.

## Diversity, equity, ethics, and inclusion [optional]

n/a.

## Data, code, and materials availability

All data, code, predicted structures, and materials used in the analysis will be available upon request.

## Supplementary Materials

Materials and Methods

Supplementary Text

Figs. S1 to S9

Tables S1 to S3

References (*32–54*)

## Materials and Methods

### Bioinformatics analyses

#### Sequence alignment and tree construction

Erv14/Cornichon sequences were collected from UniProt (*32*). Sequence alignment was performed using CLUSTALW in software package MEGA12 (*33*). The aligned sequences were subsequently used as input to construct a maximum likelihood tree by WAG (g + i) substitution model. The sequence alignment was visualised by ESPript 3.2 (*34*) and tree visualised by iTOL (*35*).

#### Structural prediction and contact analysis

Structural prediction of Erv14 in complex with different Mid2 TMH chimeras was performed using an AlphaFold2-Multimer (*36*) and ColabFold build (*37*), with 6 independent random seeds to create an ensemble of 30 predicted structures. The top five paired alignment error (PAE) matrices are shown in Fig. S2A. The overlay of 30 predictions is visualised in Fig. S2B. A contact is defined as carbon-alpha distance less than 10 Å (*38*). The mean number of contacts per Erv14 residue was calculated for the ensemble of 30 structures for each complex combination.

Structural predictions of Erv14 and Sec24 were generated using the same pipeline. For other structural predictions in the manuscript, the top ranked prediction from AlphaFold2-Multimer and ColabFold was visualized. All sequences apart from polyleucine chimeras were collected from UniProt using the canonical isoform. For Sec24 proteins, the N-terminal intrinsically disordered regions were trimmed to increase prediction accuracy. For Mid2, Axl2, Gurken, Notch, only the TMH and flanking cytoplasmic region were used as input to increase reliability. Full-length Yor1 were used as input. Due to the size of the system, Erv14-Sec24-Yor1 structure was predicted using AlphaFold3 (*39*). Input sequences and structural prediction details are reported in Table S1 and Data S1, and additional PAE matrices are summarized in Fig. S9.

Structural conservation analysis related to Fig. 1D was performed using Consurf (*40*) with default setting. Sequence logo corresponding to Fig. 3A was generated using ConservFold (*41*).

### All-atom molecular dynamics simulation and data analysis

#### Simulation Setup

Molecular dynamics (MD) simulations were organized into two groups. The first group examined the effect of bilayer thickness and included Erv14, Mid2_TMH_, and the Erv14-Mid2_TMH_ complex in single-component DRPC (1,2-dimyristoleoyl-sn-glycero-3-phosphocholine; di-14:1 PC) (thin) and DSPC (1,2-distearoyl-sn-glycero-3-phosphocholine; di-18:0 PC) (thick) bilayers. To isolate the role of bilayer thickness, we performed simulations in single-component DRPC and DSPC bilayers. DRPC has short, unsaturated acyl chains, which provide a thin, disordered membrane environment, whereas DSPC has long, saturated acyl chains, providing a thicker, more ordered membrane environment. As both lipids share the same headgroup, this setup minimizes differences in headgroup chemistry and allows us to focus primarily on how hydrophobic thickness influences Erv14, Mid2_TMH_, and the Erv14-Mid2_TMH_ complex. We therefore use these bilayers as idealized physical models of thin ER-like and thick Golgi-like membranes. The second group examined the effects of Sec24 binding and of the cytoplasmic region of Mid2, including Erv14-Sec24, Erv14-Sec24-Mid2, and Erv14-Mid2_TMH_ in a DRPC bilayer. A summary of all simulated systems is provided in Table S2.

The initial membrane-protein systems were assembled using the CHARMM-GUI Membrane Builder (*42*). For each system, the corresponding protein structure was embedded in either a DSPC or DRPC lipid bilayer, as stated in Table S2, solvated with TIP3P water. To maintain charge neutrality and mimic physiological ionic concentration, 150 mM of Na and Cl ions were added to the system. The CHARMM36 (*43*) force field family of parameters was used for all components of the system, with CHARMM36m (*44*) being used for the protein.

Following system assembly, energy minimization was performed using the steepest-descent algorithm to remove unfavourable steric contacts. Minimization was terminated when the maximum force was reduced below 500 kJ.mol^-1^nm^-1^, or after 10000 steps. Subsequently, each system was equilibrated using the standard CHARMM-GUI stepwise membrane equilibration protocol. During equilibration, positional restraints were applied to the protein backbone, protein side chains, and lipid headgroups, together with selected lipid dihedral restraints. These restraints were gradually reduced during the equilibration stages to allow relaxation of the membrane and solvent environment while maintaining the overall orientation and integrity of the membrane-embedded protein assemblies.

Equilibration was first carried out in the NVT ensemble for 1 ns using a Berendsen thermostat (*45*) at 300K with a 1 fs integration time step. This was followed by NPT equilibration for 2 ns using a C-rescale barostat (*46*) at 1 atm pressure. For membrane-containing systems, pressure coupling was applied semi-isotropically, allowing independent fluctuations of the membrane plane and bilayer normal. Production simulations were then performed in the NPT ensemble at 300 K and 1 atm pressure using the Bussi velocity rescale thermostat (*47*) and C-rescale barostat settings with semi-isotropic coupling. Long-range electrostatic interactions were computed using the particle-mesh Ewald method (*48*). Van der Waals interactions were treated with a 12 Å cutoff and a switching distance of 10 Å. The neighbor list was updated every 20 steps using a grid-based method, and hydrogen bonds were constrained using the LINCS (*49*) algorithm. The same minimization, equilibration, and production run protocols were used for all systems using GROMACS 2021.5.

#### Residue Insertion Depth Calculation

Residue insertion depth was calculated to determine how deeply each residue was positioned within the membrane. For each frame, the local positions of the two membrane leaflets were defined using lipid phosphate phosphorus atoms within 20 Å of the protein, which avoids errors from large-scale membrane bending. A best-fit plane was calculated for each leaflet, and the distance of each residue C*α* atom from the nearest leaflet plane was measured along the leaflet normal:

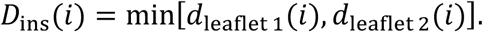

Here, *D*_ins_(*i*) reports the insertion depth of residue *i*. Negative values indicate residues located inside the hydrophobic core of the bilayer, whereas positive values indicate residues positioned outside the membrane interface.

#### Tilt Angle Calculation

Tilt angle was calculated to measure the orientation of each transmembrane helix with respect to the membrane normal. For each frame, the membrane normal was defined from the two fitted leaflet planes. The helix axis was defined based on the C*α* atoms of the selected transmembrane helical residues using singular value decomposition. The tilt angle, *θ*, was then calculated as the acute angle between the helix axis, **h**, and the membrane normal, **n***_m_*:

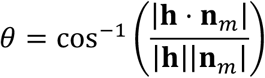

#### Bilayer Thickness Analysis

For each trajectory frame, best-fit planes were defined for the upper and lower leaflets using lipid phosphate phosphorus atoms. This was done using three selections: phosphate atoms within 1 nm of the protein, phosphate atoms within 2 nm of the protein, and all phosphate atoms in the bilayer. The first two definitions report the local bilayer thickness around the protein, whereas the full-bilayer definition reports the average membrane thickness of the system. The bilayer thickness was calculated as the distance between the upper and lower leaflet planes along the membrane normal.

#### Kink Angle Analysis

The kink angle of Mid2 was computed using HELANAL (*50*) module in MDAnalysis (*51*), using the C *α* atoms of Mid2 residues 230-236. HELANAL calculates one bend angle from two overlapping four-residue windows, residues 230-233 and 233-236, which define the two local helix directions. The kink angle was calculated as

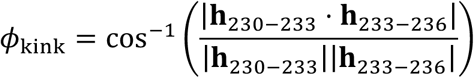

where **h**_230−233_and **h**_233−236_are the local helix direction vectors calculated from the C*α* atoms of residues 230-233 and 233-236, respectively.

#### Root Mean Square Deviation (RMSD)

Root-mean-square deviation (RMSD) measures the overall structural deviation of a protein from a reference conformation over time, providing insight into the extent of conformational difference from the reference structure. For all Erv14-related comparisons, the Erv14-Mid2-Sec24 structure was used as the common reference structure. Trajectory frames were aligned by fitting the helical region of Erv14 to the corresponding region in the reference structure. The RMSD was then calculated as

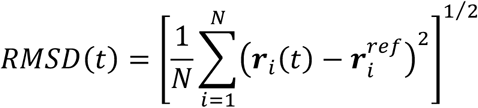

where N is the number of atoms included in the calculation, ***r****_i_*(*t*) is the aligned position of the atom *i* at time *t*, and 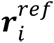 is the corresponding position in the reference structure. This procedure was used to compare Erv14 loop RMSD across Erv14, Erv14-Mid2, Erv14-Sec24, Erv14-cytMid2-Sec24, and Erv14-cytMid2 systems.

For the calculation of the RMSD of the cytoplasmic region of Mid2, trajectory frames were aligned using the transmembrane region of Mid2.

#### Root Mean Square Fluctuation (RMSF)

Root-mean-square fluctuation (RMSF) analysis was used to quantify residue-level flexibility. For each system, an average protein structure was generated from the aligned trajectory with respect to the first frame, and the RMSF was calculated as

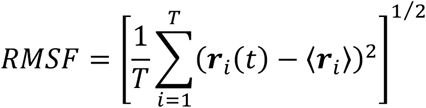

where *T* is the number of trajectory frames, **r***_i_*(*t*) is the aligned position of the atom or residue *i* at time *t*, and ⟨**r***_i_*⟩ is its average position over the trajectory.

For the calculation of the RMSF of the loop regions of Erv14, frames were fitted using the helical region of Erv14, and fluctuations were calculated relative to the average Erv14 structure. For the calculation of the RMSF of the cytoplasmic region of Mid2, frames were fitted using the transmembrane region of Mid2, and fluctuations were calculated relative to the corresponding average structure.

### Yeast molecular genetics and phenotype analyses

Yeast strains and plasmids are summarized in Table S3. Standard yeast transformation protocols using PEG-LiAc were applied to introduce plasmids in yeast cells (*52*) with corresponding auxotrophic marker selections. Site-directed mutagenesis was performed using a QuikChange Lightening kit according to the manufacturer’s protocol. DNA primers were ordered from IDT DNA. Mutations were confirmed by whole-plasmid sequencing. Erv14 mutants were tested in the parental strain: BY4741 *erv14:KanMX, erv15:NatMX, mid2-GFP::HisMX*. This strain was created by chromosomal tagging of the *MID2* locus with a GFP::HisMX cassette using the Longtine PCR method. Mid2 mutants were analyzed in the parental strain: YPH499, *sec24::TRP*, *pRS316-SEC24* (URA).

#### Immunoblot and quantification

For collection of samples for immunoblots, cells were grown at 30°C to mid-log phase (OD_600_ 0.5–0.65). A total of 5.5 OD cells were collected for each sample by centrifugation. Cell pellets were either lysed immediately or stored at −20 °C and lysed within a week. Cell pellets were lysed as previously described (*18*). Briefly, the sample was resuspended in 100µL water at room temperature. Equal volume of 0.2M NaOH was added to the suspension, and the mixture was incubated for 5 min at room temperature, then placed on ice. Cells were collected by centrifugation (12.000 rpm, 5min, 4°C) and resuspended in 35µL 3xSB buffer (0.187 M Tris pH 6.8, 30% [v/v] glycerol, 6% [w/v] SDS, 0.1% [w/v] bromophenol-blue and 0.2 M dithiothreitol [DTT]) and heated for 10 min at 55°C. Lysates were cleared by centrifugation (12.000 rpm, 5min, 4°C), 30uL of supernatant collected, and total protein concentration was estimated using a quick Bradford assay (Bio-Rad) for normalization of protein loading.

Lysates were separated on 4–12% NuPAGE Bis–Tris gels, transferred to PVDF membrane and immunoblotted with a mouse anti-HA antibody (diluted 1:10,000) and HRP-conjugated goat anti mouse secondary antibody (diluted 1:5,000) to detect Erv14-3xHA. Mid2-GFP was detected using mouse monoclonal anti-GFP antibody (Roche; 1:1,000 dilution), followed by HRP-conjugated goat anti mouse secondary antibody diluted 1:5,000.

The blots were imaged using a Bio-Rad GelDoc system in manual exposure mode, following incubation with enhanced chemoluminescence substrate (Pierce). Images were subsequently quantified by densitometry using ImageLab (BioRad) software.

#### Data analysis of immunoblots

To quantify effects of mutation of the H1-H4 interface in Fig. 1, Mid2-GFP maturation was measured independently in at least 4 biological replicates. Each immunoblot contained one lane of wild type Erv14-3xHA, and one lane of empty vector, for normalization. The pixel intensity of ER from, [*ER form*]*_mut_*, and Golgi form, [*Golgi form*] for each lane was measured, and the ER fraction calculated as

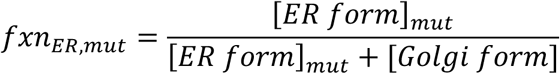

The logarithm of the ER fraction of each mutant normalized to the wild type on the same blot is then reported as fraction of enrichment for each mutant, *log*_2_([*mut*]/[*WT*]), in Fig.1, where:

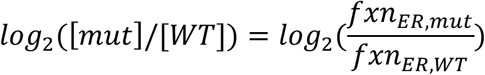

Data for each mutant was then log-transformed [*log*_2_([*mut*]/[*WT*])] to compare to WT, then subject to a one-sample t-test against null hypothesis mean zero followed by Benjamini– Hochberg correction with false discovery rate (FDR) at 0.05, to determine the significance of mutational effect. The logarithm of adjusted p value is reported in Fig.1.

For loop mutants related to Fig. 4 and Fig. S3C, the ER fraction was calculated as described above. The statistical significance of mutational effect was determined t-test against null hypothesis mean zero, followed by Holm Sidak correction. Statistical analyses were performed using Prism 11.

### Drosophila transfection and phenotypic assessment

#### Generation of Cornichon control and mutant lines

Fly lines are listed in Table S3. We synthesized the same genomic region as used in a previously published Cornichon rescue construct (2L:16311560-16309030, *Drosophila melanogaster* Release 6.67) (*7*), added a C-terminal 3xHA tag and applied the following additional mutations: none (Cni^WT^::3xHA); ACA209-211GCC, ATA221-223GCC (Cni^I25A,I29A^::3xHA); ACCACGGTCCTGAAA630-644GCCGCTGCGGCCGCT (Cni^104-108A^::3xHA); ATATACC720-725GCCGCC (Cni^I134A,Y135A^::3xHA) where nucleotide positions refer to the Cni-RA transcript (FlyBase: FBtr0080848). This was then inserted into a plasmid with a miniWhite selection marker and an attB site. Transgenic lines were generated by site-directed transgenesis (https://www.flyfacility.gen.cam.ac.uk/Services/Microinjectionservice) into attP2.

#### Immunostaining of Drosophila egg chambers

At room temperature, Drosophila ovaries from adults 24h after pupal hatching were dissected into individual ovarioles in phosphate buffered saline (PBS), fixed in 4% Formaldehyde (Sigma) for 12 minutes, rinsed twice in PBS, washed 15 minutes in PBS and permeabilised for 2 hours in PBS-Triton 0.05% (PBT). Ovarioles were then incubated overnight at 4°C in a solution of Mouse-anti-Gurken antibodies (DSHB) diluted 1/200 (Gurken staining) or Rabbit-anti-HA antibodies diluted 1/200 (Cornichon::HA staining) in PBT. The next day, at room temperature, ovarioles were rinsed three times in PBT, washed 2 hours in PBT, incubated 1 hour in a solution of Goat-anti-Mouse-Alexa647 secondary antibodies (Thermo Fisher) 1/1000 and Phalloidin-Alexa594 1/10000 in PBT (Gurken staining) or Donkey-anti-Rabbit-Alexa594 secondary antibodies (Thermo Fisher) 1/1000 (Cornichon::HA staining), rinsed twice in PBT, washed 45 minutes in PBT, rinsed twice in PBS, rinsed once in 50% glycerol (Sigma), and finally mounted between glass and coverslip in Vectashield Plus (Vector Laboratories H-1900). Stage 9 and 10A (for Gurken staining) or stage 2-4 (Cornichon::HA staining) egg chambers were imaged on a LEICA SP8 Stellaris confocal microscope equipped with an 86x water immersion objective (NA 1.20).

#### Fluorescence microscopy processing and analysis

All images were processed and analysed using ImageJ (*53*). A 2D gaussian blur with a sigma of 0.8 was applied to all fixed fly tissues images displayed in Figures, but we performed intensity measurements on images without a gaussian blur. Gurken levels at the plasma membrane were measured as the average intensity in a 12 µm-long band overlapping with the Phalloidin signal, wherever the Gurken signal was the brightest (typically close to the oocyte nucleus in stage 9 egg chambers and more posteriorly in stage 10A egg chambers). We measured the signal within the nucleus to estimate the autofluorescence, which we subtracted from the Gurken signal at the plasma membrane. We then normalised these values to the average signal in controls. As optical sections gather more signal from the plasma membrane when it is orthogonal rather than parallel to the plane of imaging, we independently normalised all “orthogonal” measurements to “orthogonal” controls, and all “parallel” measurements to “parallel” controls before pooling the data. To quantify Cni enrichment at the ERES, for each egg chamber, nurse cells were manually segmented (the signal from follicle cells and the oocyte was erased), Sec16-positive punctae were automatically segmented based on their intensity, the average Cni intensity was individually measured in each Sec16-positive puncta, and the measured values were divided by the average Cni signal within the entire Cni-positive compartment after background subtraction. Statistical significance for each pair was determined by ordinary one-way ANOVA followed by post-hoc Dunnett’s (Fig. 3C) or Tukey’s (Fig. 3F) correction, for correcting multiple comparisons.

#### Drosophila wings analysis

One wing was analysed per animal. Right wings were cut from adult flies, mounted dry between slide and coverslip, and imaged on a Leica MSV269 Stereo Microscope. Images were processed and analysed using ImageJ (*53*). The ImageJ “Subtract Background” function was applied with the following settings: Rolling ball radius: 900 px; Light background enabled; Create background disabled; Sliding paraboloid enabled; Smoothing enabled. For analysis only (not displayed in Figures), in order to better visualize the exact point where the L5 vein is interrupted, we then applied a bandpass filter (Filter large structure down to 1 px; Filter small structure down to 15 px; Suppress strips: None; Tolerance of direction: 0; Autoscale after Filtering; Saturate image when autoscaling) to filter out the small epidermal hairs. We calculated a “L5 interruption percentage” indicating which percentage of the section of the L5 vein from its intersection with the posterior crossvein (p-cv) to the margin is missing (see Fig. 3D for a graphical explanation). Statistical significance for each pairwise comparison was determined by Kruskal-Wallis test followed by post-hoc Dunn’s test to correct for multiple comparisons.

### General consideration in considering statistical significance

Our multidisciplinary approach yielded data from different biological backgrounds, sampling design, and quantification procedures. We therefore chose appropriate statistical tests based on data normalcy, distribution, continuity, independency in sampling, and other factors that may affect the null hypothesis for each individual design. For figures with multiple statistic tests, correction methods that are suitable for specific experimental designs were chosen to improve accuracy of statistic inference.

---

**Fig. S1.**
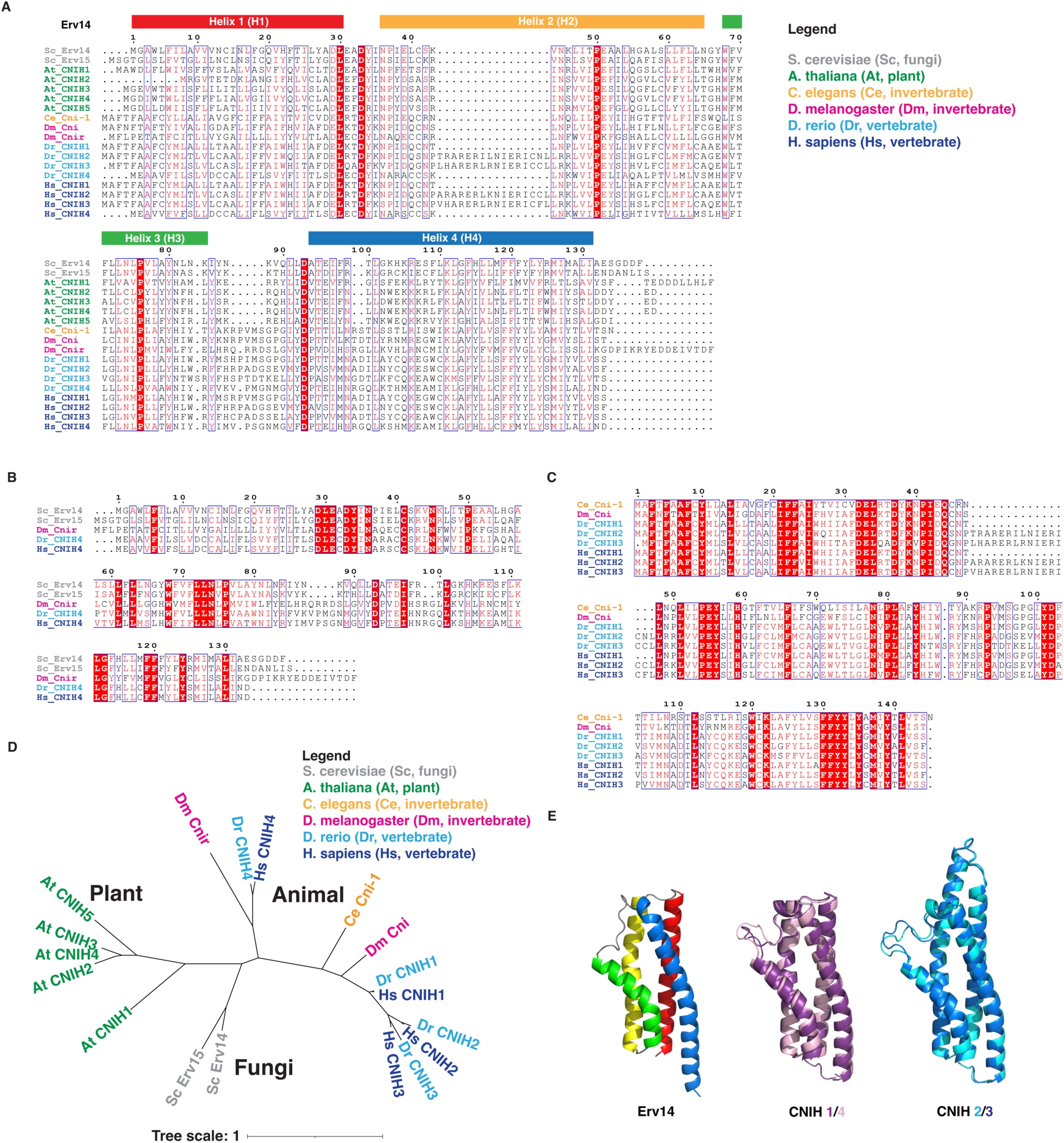
Cornichon proteins are highly conserved in eukaryotes. (A) Sequence alignment for all cornichon homologs from model organisms. (B) Sequence alignment for Erv14-CNIH4 orthologs from model organisms. (C) Sequence alignment for animal-specific cornichon orthologs from model organisms. (D) Evolutionary tree of cornichon homologs across different model organisms, showing cornichon homologs widely exist in eukaryotes. Tree scale bar stands for one mutation per position. (E) AlphaFold2 predictions for yeast Erv14 (left), human CNIH 1/4 (middle), and human CNIH 2/3 (right).

**Fig. S2.**
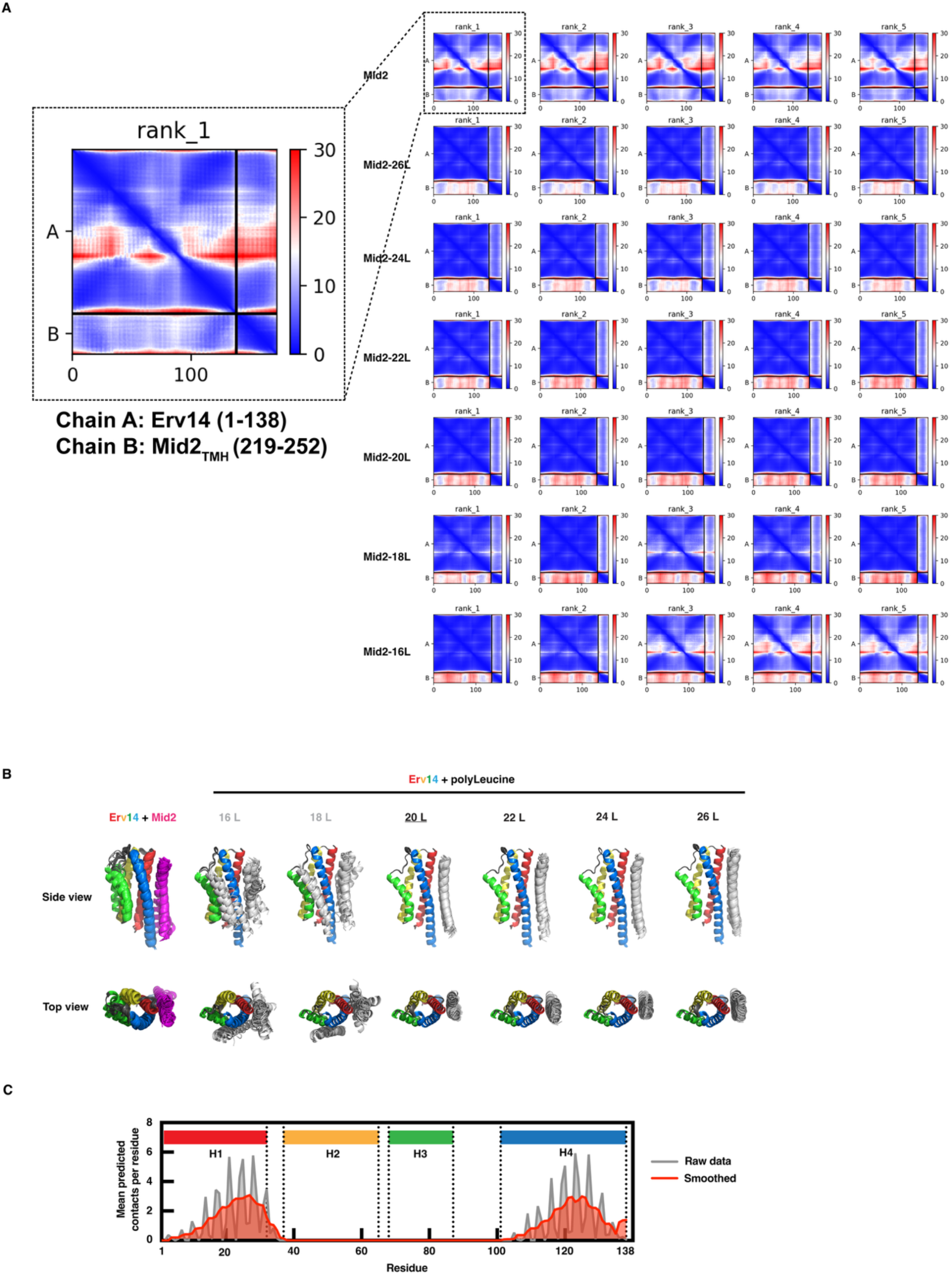
AlphaFold2 predictions of Erv14-Mid2 chimera complexes. (A) Predicted alignment error (PAE) matrices for top five ranked models across wild type Mid2 and six Mid2 chimeras. Apart from the 16L chimera which has relatively high PAE values for H1-H4 interface in low-ranked models, all other predictions resulted in robust interface between Mid2 TMH and Erv14 H1-H4 interface. In the predictions, chain A represents Erv14, chain B represents various Mid2 chimeras. (B) Overlay of 30 predicted conformations of complexes between Erv14 and Mid2 chimeras, corresponding to Fig. 1A. (C) Contact probability reflected by mean predicted contacts between Erv14 and wild type Mid2 per Erv14 residue, plotted as raw number of predicted contacts (grey) and smoothed data (red) to reflect the trend. The high probability of contacts locates within H1 and H4.

**Fig. S3.**
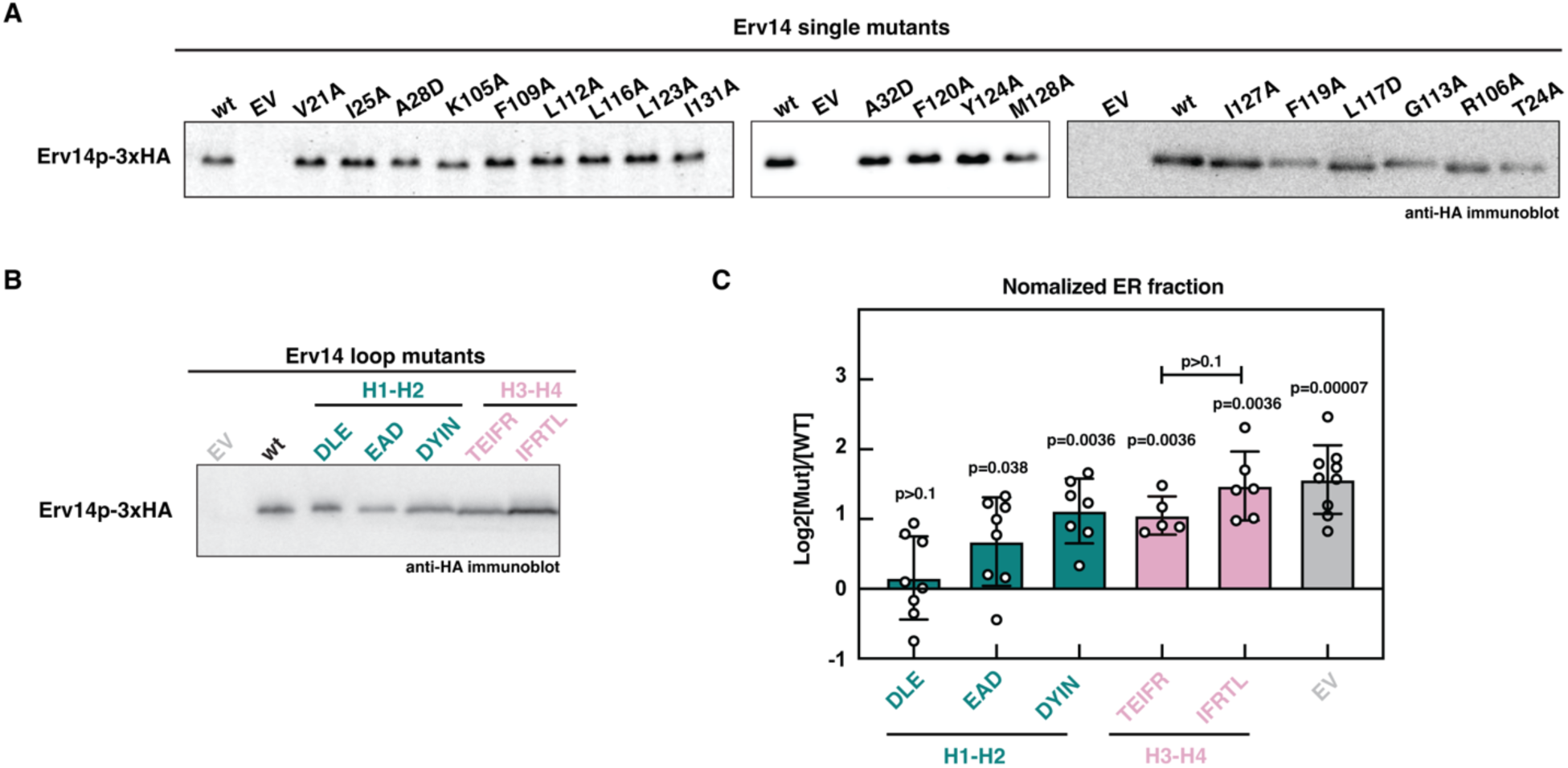
Mutant protein stability and Mid2 trafficking quantification. (A) Steady-state expression levels of Erv14 single mutants used in Fig. 1. (B) Steady state expression levels of Erv14 loop mutants used in Fig. 4. (C) Quantification of immunoblots related to Fig. 4B. Fisher’s LSD test for ANOVA was used. The significance of each column against wild type (calculated as 0, not shown) is shown on top of each bar. (One sample t-test against null hypothesis mean zero, followed by Holm Sidak correction, n = 5-8). No significant difference was observed between TEIFR and IFRTL signal mutations in H3-H4 loop (Welch’s t-test).

**Fig. S4.**
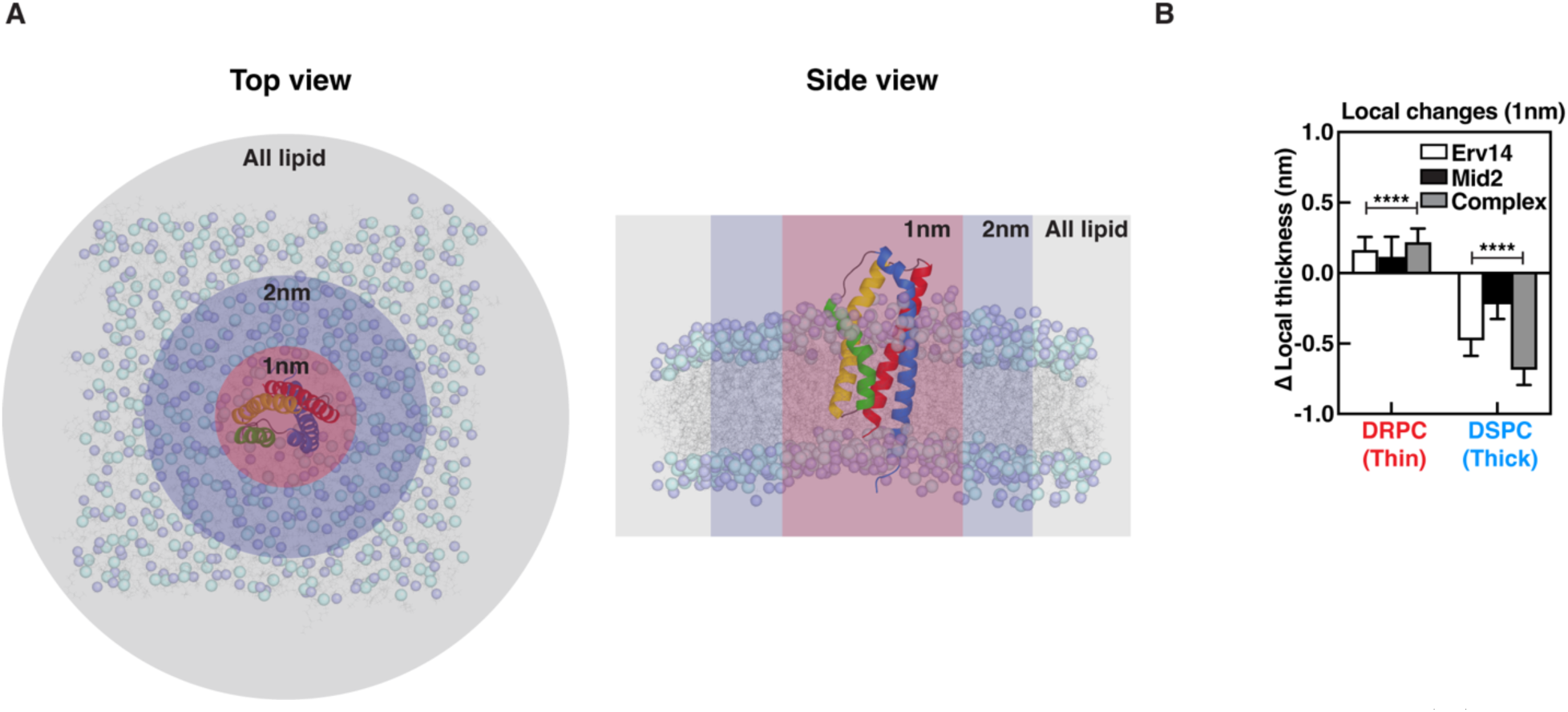
Scheme and calculation of bilayer thickness modulation in MD simulation. (A) Top view (left) and side view (right) of a bilayer with Erv14 embedded. The mean bilayer thickness across all lipids (grey), 2 nm from the surface of Erv14 (blue), and 1 nm from the surface of Erv14 (red) were calculated and reported in Figure 2. Cyan and violet color bead represents the P atom and N atom in the phosphocholine headgroup respectively. (B) Comparison of change in local bilayer thickness at 1nm from centre of mass of Erv14 alone (white), Mid2 alone (black), or Erv14-Mid2 (grey). Binding of Mid2 to Erv14 further amplified the modulation of bilayer thickness by Erv14 (multiple unpaired t-test, n = 8000; ****: p<0.0001).

**Fig. S5.**
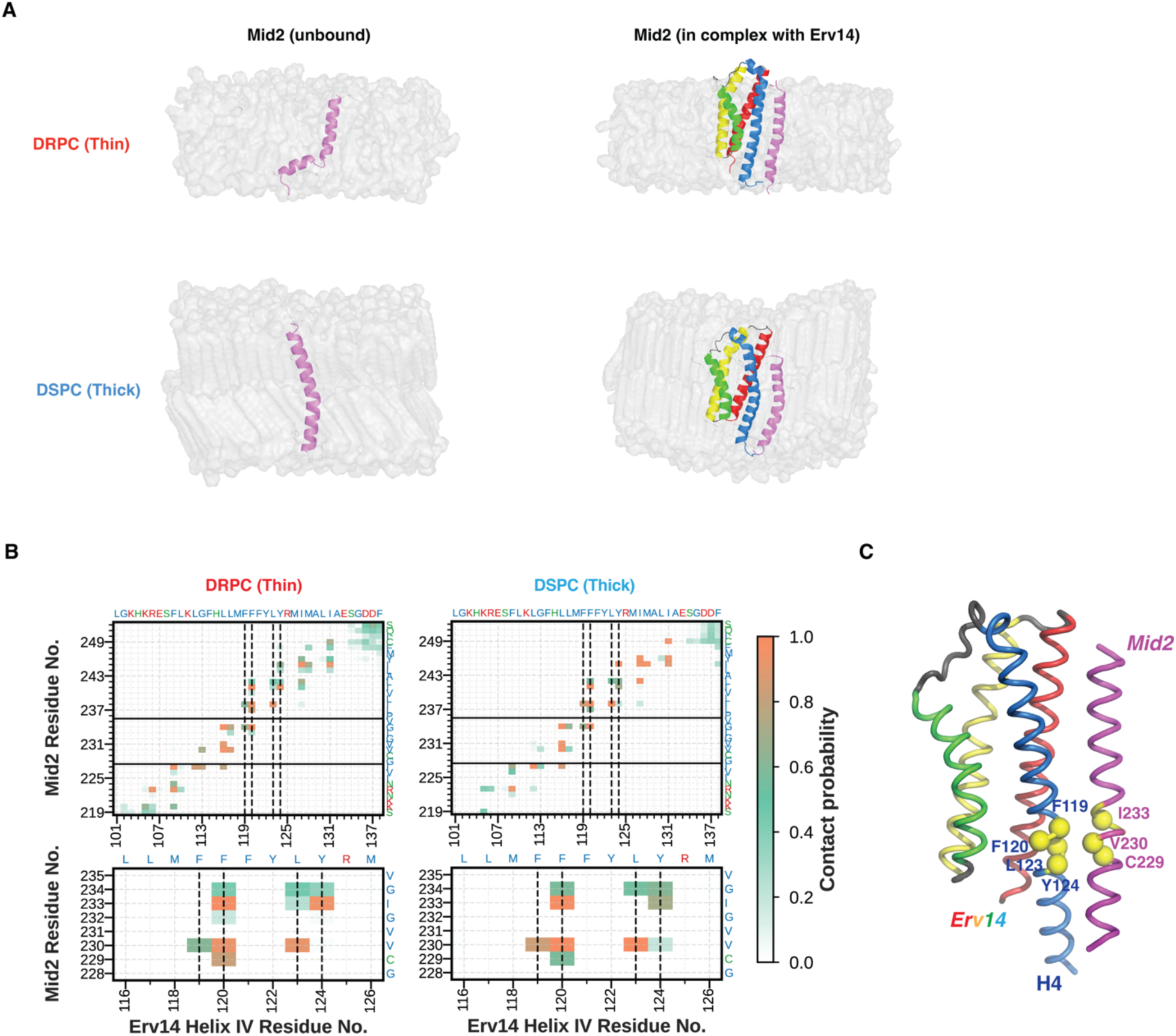
Erv14 rectifies the Mid2_TMH_ kink caused by hydrophobic mismatching. (A) Snapshots of representative conformations of Mid2_TMH_ (magenta) in DRPC/DSPC with or without Erv14 (rainbow). (B) Contact frequency calculated from 50,000 frames of MD simulation between Mid2_TMH_ and H4 of Erv14. Of all residues in Erv14 H4, residues F119, F120, L123, Y124 (marked by dashed lines) have the most contact with Mid2_TMH_ residues in the helix kink. (C) Cartoon illustration of Erv14 H4 (blue) -Mid2 (magenta) kink contact in Fig. S5B. Residues with bulky hydrophobic side chains are highlighted by yellow spheres.

**Fig. S6.**
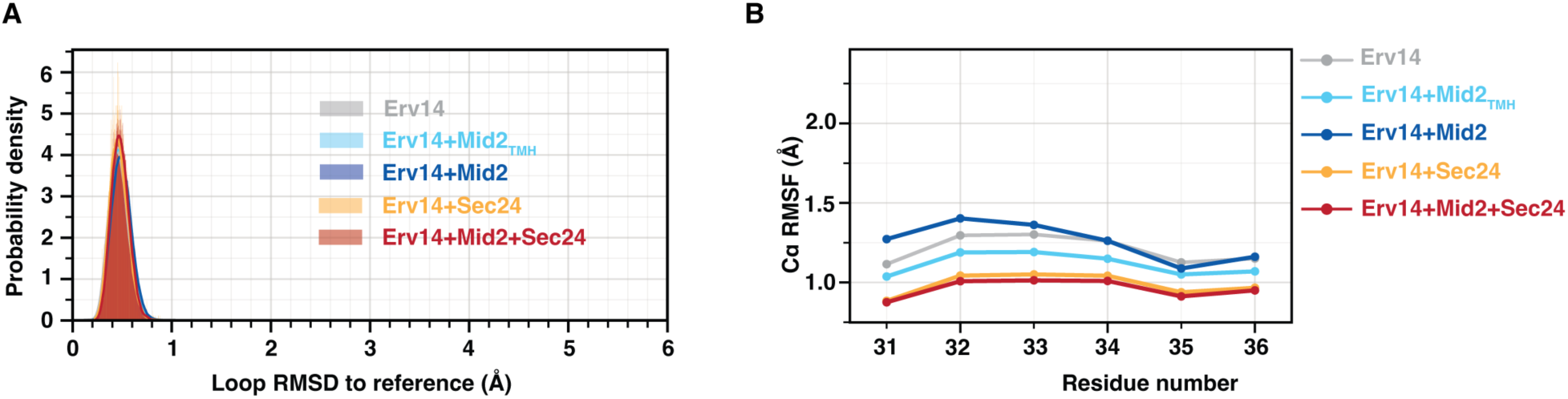
RMSD and RMSF analysis related to H1-H2 loop of Erv14. (A) RMSD distribution of H1-H2 loop in Erv14 for each simulation relative to the reference structure of Erv14-Mid2-Sec24. (B) Residue-specific RMSF data across H1-H2 loop in Erv14.

**Fig. S7.**
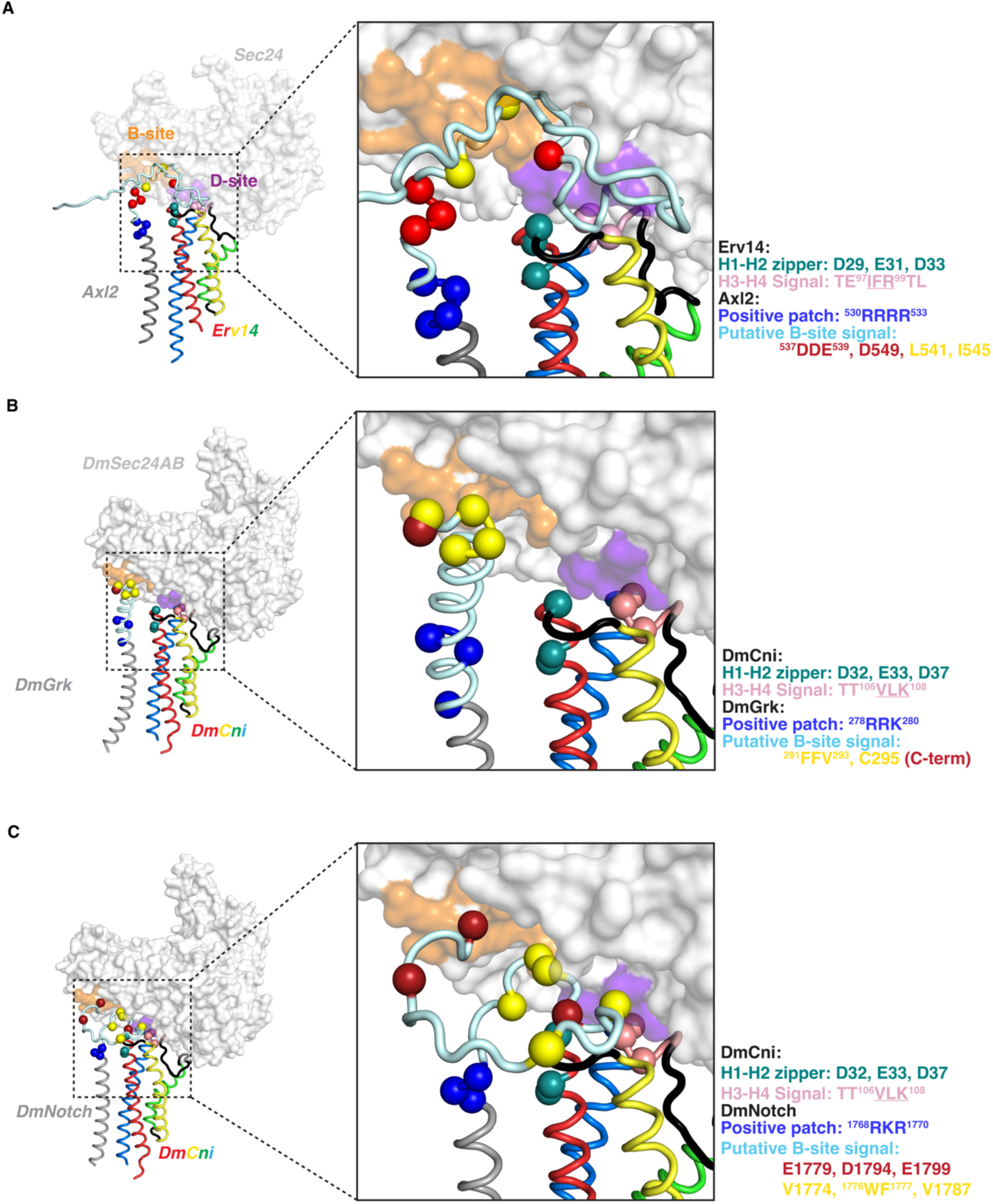
Conservation of HEIST motifs in Erv14/Cornichon clients. Structural predictions for (A) Axl2p-Erv14-Sec24. (B) Gurken-CNI-Sec24AB. (C) Notch-CNI-Sec24AB. In each structure, Sec24 orthologs are colored white, with conserved B-site and D-site shaded orange and purple respectively. Cornichon homologs are modeled as rainbow tubes, matching coloring in the main text (H1: red, H2: yellow, H3: green, H4: blue). Key structural motifs involved in HEIST are colored in the same convention as main text, where the putative B-site signals on cytoplasmic loops region of each cargo were shown as spheres and colored accordingly.

**Fig. S8.**
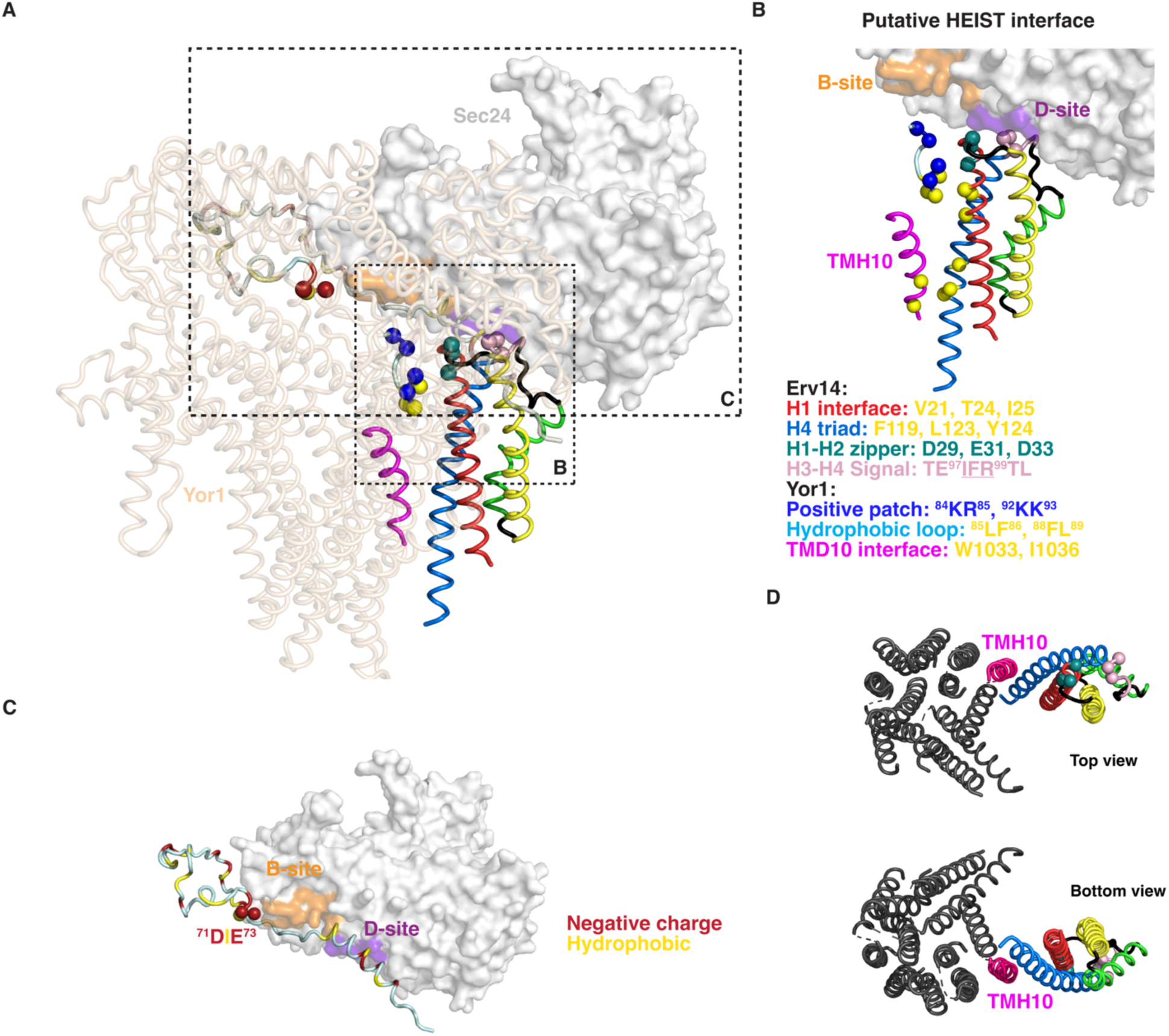
Conserved HEIST motifs in a polytopic Erv14 client. (A) Predicted structure of Yor1-Erv14-Sec24. (B) The HEIST interface predicted for Yor1 consists of Yor1 TMH10 (magenta) interacting with Erv14-H4 (blue), plus a hydrophobic interaction at the N-terminus of H1 (red tube, yellow spheres) that engages with hydrophobic residues in a cytoplasmic loop of Yor1 (cyan tube, yellow spheres) that are flanked by positive charge patches (cyan tube, blue spheres) that may engage the Erv14 H1-H2 zipper (red tube, green spheres). Key residues are described in the figure. (C) The Sec24 B-site (orange) is in proximity to an N-terminal disordered region of Yor1 that carries both acidic and hydrophobic residues, including the previously characterized B-site signal ^71^DIE^73^ (red spheres). (D) Illustration of helix engagement from top/bottom views between Yor1-TMH10 (magenta) and Erv14 (rainbow).

**Fig. S9.**
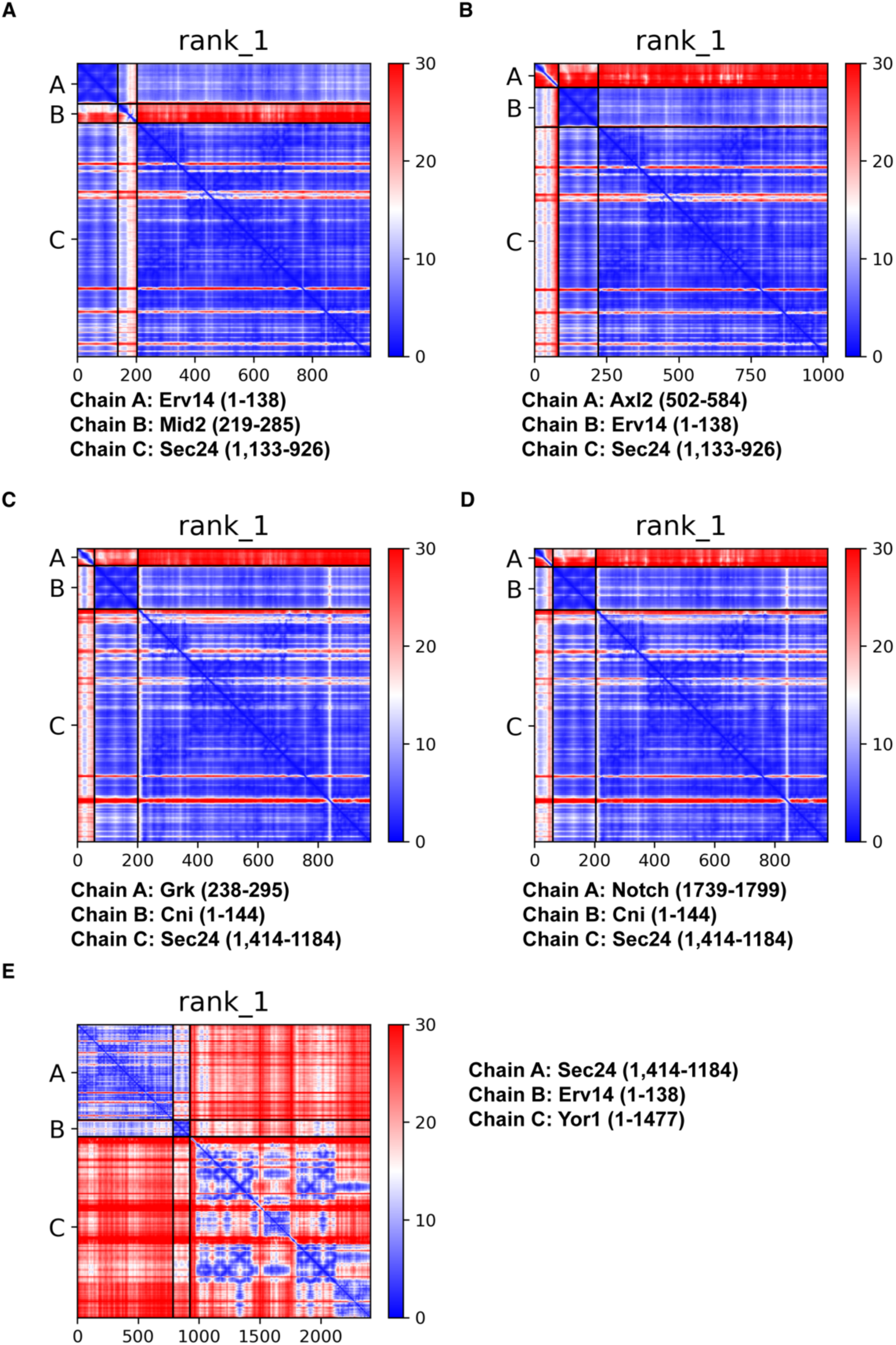
Predicted alignment error (PAE) matrices for structural predictions of heterotrimer complexes. PAE matrices for the best ranked model related to (A) Erv14-Mid2-Sec24, (B) Erv14-Axl2-Sec24, (C) Cni-Grk-Sec24, (D) Cni-Notch-Sec24, (E) Erv14-Yor1-Sec24.

**Table S1.** Details of predicted structures used in this work.

| Entry | Chain 1 | Chain 2 | Chain 3 | Appearance | Algorithm/<br>Resource |
| --- | --- | --- | --- | --- | --- |
| **Erv14-Mid2<sub>TMH</sub>** | Erv14 [P53173]<br>(1-138) | Mid2 [P36027]<br>(219-252) |  | Fig.1, Fig. S2 | AlphaFold2-multimer-v3 |
| **Erv14-Mid216L** | Erv14 [P53173]<br>(1-138) | Mid2-16L<br>chimera |  | Fig.1, Fig. S2 | AlphaFold2-multimer-v3 |
| **Erv14-Mid218L** | Erv14 [P53173]<br>(1-138) | Mid2-16L<br>chimera |  | Fig.1, Fig. S2 | AlphaFold2-multimer-v3 |
| **Erv14-Mid220L** | Erv14 [P53173]<br>(1-138) | Mid2-16L<br>chimera |  | Fig.1, Fig. S2 | AlphaFold2-multimer-v3 |
| **Erv14-Mid222L** | Erv14 [P53173]<br>(1-138) | Mid2-16L<br>chimera |  | Fig.1, Fig. S2 | AlphaFold2-multimer-v3 |
| **Erv14-Mid224L** | Erv14 [P53173]<br>(1-138) | Mid2-16L<br>chimera |  | Fig.1, Fig. S2 | AlphaFold2-multimer-v3 |
| **Erv14-Mid226L** | Erv14 [P53173]<br>(1-138) | Mid2-16L<br>chimera |  | Fig.1, Fig. S2 | AlphaFold2-multimer-v3 |
| **Erv14-Mid2** | Erv14 [P53173]<br>(1-138) | Mid2 [P36027]<br>(219-285) |  | Used in MD<br>simulation | AlphaFold2-multimer-v3 |
| **Erv14-Sec24** | Erv14 [P53173]<br>(1-138) | Sec24 [P40482]<br>(1,133-926) |  | Used in MD<br>simulation | AlphaFold2-multimer-v3 |
| **Erv14-Mid2-Sec24** | Erv14 [P53173]<br>(1-138) | Mid2 [P36027]<br>(219-285) | Sec24 [P40482]<br>(1,133-926) | Fig. 4 | AlphaFold2-multimer-v3 |
| **Erv14-Axl2-Sec24** | Erv14 [P53173]<br>(1-138) | Axl2 [P38928]<br>(501-584) | Sec24 [P40482]<br>(1,133-926) | Fig. S7 | AlphaFold2-multimer-v3 |
| **Cni-Grk<sub>TMH</sub>-**<br>**Sec24AB** | Cni [P49858]<br>(1-144) | Grk [P42287]<br>(238-295) | Sec24AB<br>[A1Z813]<br>(1,414-1184) | Fig. S7 | AlphaFold2-multimer-v3 |
| **Cni-Notch<sub>TMH</sub>-**<br>**Sec24AB** | Cni [P49858]<br>(1-144) | Notch [P07207]<br>(1739-1899) | Sec24AB<br>[A1Z813]<br>(1,414-1184) | Fig. S7 | AlphaFold2-multimer-v3 |
| **Erv14-Yor1-Sec24** | Erv14 [P53173]<br>(1-138) | Yor1 [P53049]<br>(1-1477) | Sec24 [P40482]<br>(1,133-926) | Fig. S8 | AlphaFold3 |
| **Erv14** | Erv14 [P53173]<br>(1-138) |  |  | Fig. S1 | AlphaFoldDB |
| **CNIH1** | CNIH1<br>[O95406]<br>(1-144) |  |  | Fig. S1 | AlphaFoldDB |
| **CNIH2** | CNIH2<br>[Q6PI25]<br>(1-160) |  |  | Fig. S1 | AlphaFoldDB |
| **CNIH3** | CNIH3<br>[Q8TBE1]<br>(1-160) |  |  | Fig. S1 | AlphaFoldDB |
| **CNIH4** | CNIH4<br>[Q9P003]<br>(1-139) |  |  | Fig. S1 | AlphaFoldDB |

**Table S2.** Summary of molecular dynamics simulation systems.

| **System** | **Bilayers** | **# of replicas** | **Production run length ($\mu s$)** |
| --- | --- | --- | --- |
| **Erv14** | DSPC and DRPC | 2 | 1 |
| **Mid2<sub>TMH</sub><sup>a</sup>** | DSPC and DRPC | 2 | 1 |
| **Erv14-Mid2<sub>TMH</sub><sup>a</sup>** | DSPC and DRPC | 2 | 1 |
| **Erv14-Sec24<sup>b</sup>** | DRPC | 3 | 1 |
| **Er14-Mid2<sup>c</sup>** | DRPC | 3 | 1 |
| **Erv14-Mid2-Sec24** | DRPC | 3 | 1 |

**a.** Mid2<sub>TMH</sub> denotes residues 219–252, corresponding to the transmembrane region alone.
**b.** Sec24 denotes yeast Sec24 with disordered residues (2-132) removed.
**c.** Mid2 denotes residues 219–285, which includes both the transmembrane region and the cytosolic extension of Mid2.

**Table S3.**
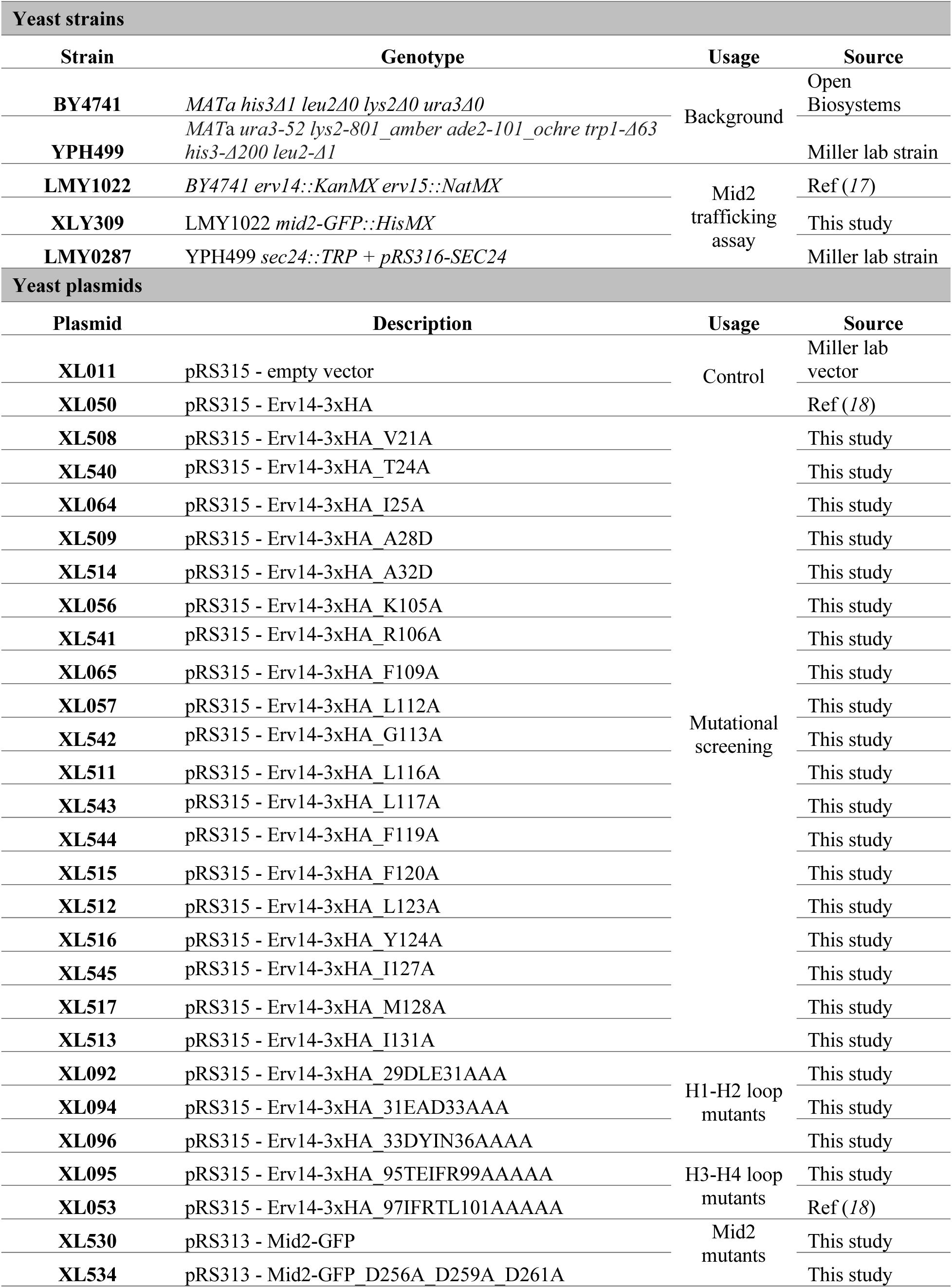
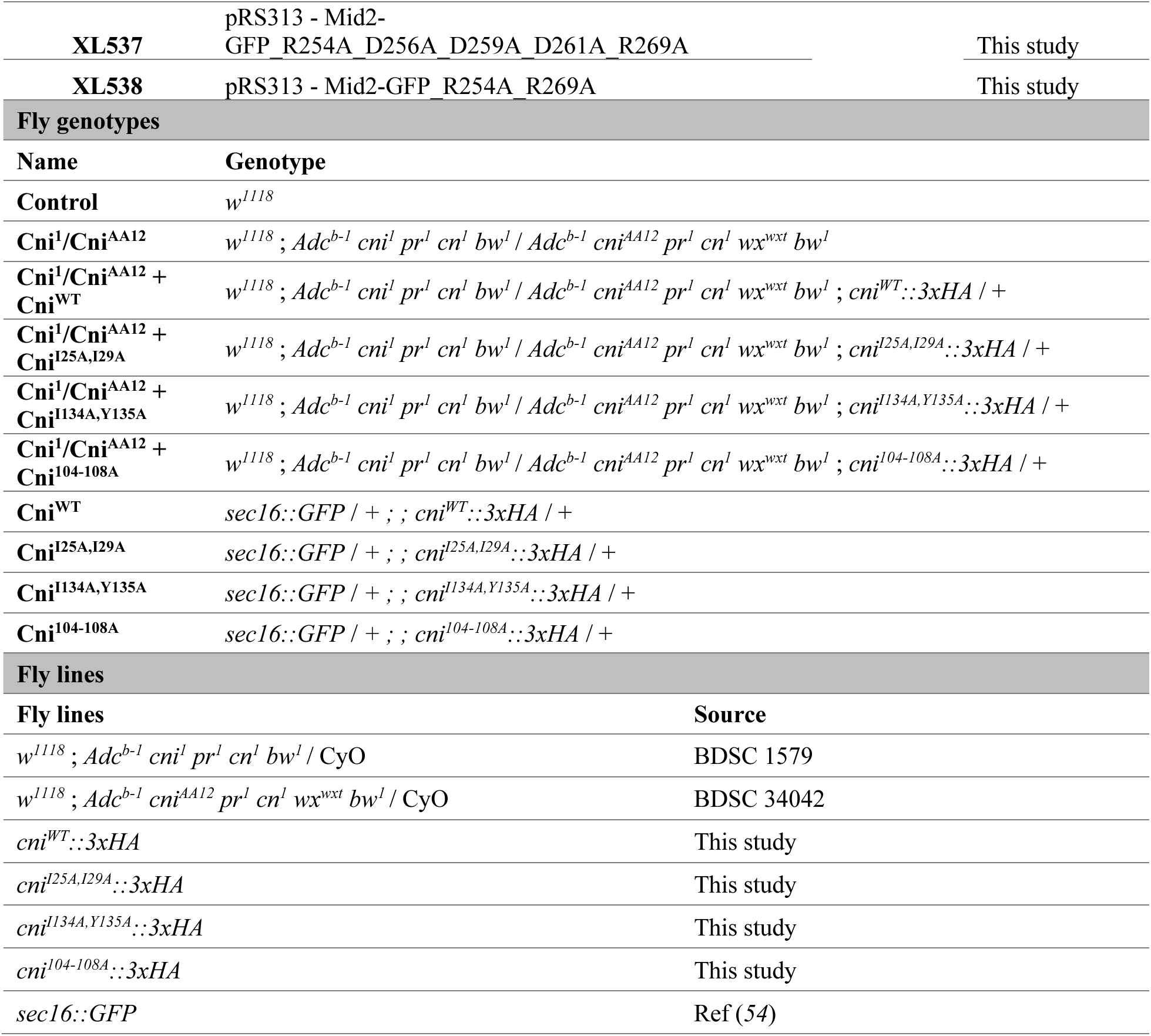
Yeast strains, fly strains, plasmids, and other resources used in this work.

